# A novel and conserved cell wall enzyme that can substitute for the Lipid II synthase MurG

**DOI:** 10.1101/2020.10.12.336396

**Authors:** L. Zhang, K. Ramijan, V.J. Carrión, L.T van der Aart, J. Willemse, G.P. van Wezel, D. Claessen

## Abstract

The cell wall is a stress-bearing structure and a unifying trait in bacteria. Without exception, synthesis of the cell wall involves formation of the precursor molecule Lipid II by the activity of the essential biosynthetic enzyme MurG, which is encoded in the division and cell wall synthesis (*dcw*) gene cluster. Here we present the discovery of a novel cell wall enzyme that can substitute for MurG. A mutant of *Kitasatospora viridifaciens* lacking a significant part of the *dcw* cluster including *murG* surprisingly produced Lipid II and wild-type peptidoglycan. Genomic analysis identified a distant *murG* paralogue, which encodes a putative enzyme that shares only around 31% aa sequence identity with MurG. We show that this enzyme can replace the canonical MurG, and we therefore designated it MurG2. Orthologues of *murG2* are present in 38% of all genomes of *Kitasatosporae* and members of the sister genus *Streptomyces*. CRISPRi experiments showed that *K. viridifaciens murG2* can also functionally replace *murG* in *Streptomyces coelicolor*, thus validating its bioactivity and demonstrating that it is active in multiple genera. Altogether, these results identify MurG2 as a *bona fide* Lipid II synthase, thus demonstrating plasticity in cell wall synthesis.

## INTRODUCTION

Bacteria are surrounded by a cell wall, which is a highly dynamic structure that provides cellular protection and dictates cell shape. A major component of the cell wall is peptidoglycan (PG), which is widely conserved in the bacterial domain. Its biosynthesis has been studied for many decades, reinforced by the notion that many successful antibiotics target important steps in this pathway. The first steps of the PG synthesis pathway occur in the cytoplasm, where the peptidoglycan precursor UDP-MurNAc-pentapeptide is synthesized by the consecutive activity of a number of so-called Mur enzymes (MurA-F)^1^. Next, this pentapeptide precursor is linked to undecaprenyl phosphate (or bactoprenol) residing in the plasma membrane by MurX (or MraY), yielding Lipid I. UDP-N-acetylglucosamine--N-acetylmuramyl-(pentapeptide) pyrophosphoryl-undecaprenol N-acetylglucosamine transferase (MurG) then adds the sugar nucleotide UDP-GlcNAc to Lipid I to form Lipid II, which is the complete PG subunit that is flipped to the external side of the membrane. Among the candidates to mediate this flipping, FtsW, MurJ and AmJ have been proposed^2–4^. Following flipping to the exterior of the cell, the PG subunit is then used to synthesize glycan strands by the activity of transglycosylases, after which these strands are cross-linked using transpeptidases ^5–8^. Many of the genes required for the biosynthesis of PG and for cell division are located in the so-called *dcw* gene cluster (for division and cell wall synthesis ^9,10^ (see Fig. S1). The content and organization of the *dcw* cluster are generally conserved among species with similar morphologies, indicating a putative role in bacterial cell shape ^11^.

Members of the *Streptomycetaceae* within the Actinobacteria are filamentous Gram-positive soil bacteria that have a complex multicellular life cycle ^12,13^. The best-studied genus is *Streptomyces*, which is industrially highly relevant as it produces over half of all known antibiotics used in the clinic, and many other bioactive compound with clinical or agricultural application^14,15^. The life cycle of streptomycetes starts with the germination of a spore, and the arising vegetative hyphae grow out via tip extension and branching to form a dense network called the vegetative mycelium. The vegetative mycelium consists of long multinucleated syncytial cells separated by widely spaced crosswalls ^16,17^. The reproductive phase is initiated by the formation of an aerial mycelium, whereby the vegetative hyphae are cannibalized as a substrate ^18,19^. The aerial hyphae then differentiate into chains of unigenomic spores. During sporulation, the conserved cell division protein FtsZ initially assembles in long filaments in the aerial hyphae, then as regular foci, to finally form a ladder of Z-rings ^20^. Eventually, cytokinesis results in spore formation, following a complex process of coordinated cell division and DNA segregation^21,22^.

Comparison between *Bacillus* and *Streptomyces* shows that some cell division-related proteins have evolved different functionalities between Firmicutes and Actinobacteria. An example of such a divergent function is exemplified by DivIVA: in *Bacillus subtilis* this protein is involved in selection of the division site by preventing polar accumulation of FtsZ ^23^, while DivIVA in Actinobacteria plays an essential role in polar growth ^24^. Thus, *divIVA* cannot be deleted in Actinobacteria while it is dispensable in *B. subtilis*. Conversely, many cell division genes, including *ftsZ*, can be deleted in Actinobacteria while being essential for unicellular microbes. This makes Actinobacteria intriguing model systems for the study of cell division and growth ^21,25^. It is also worth noticing the Streptomycetes have a complex cytoskeleton, with many intermediate filament-like proteins required for hyphal integrity ^26–29^.

Besides the genus *Streptomyces*, the family of *Streptomycetaceae* also encompasses the genera *Kitasatospora* and *Streptacidiphilus*. While highly similar in growth and development, *Kitasatospora* is distinct from *Streptomyces* ^30,31^. We recently described that *Kitasatospora viridifaciens* releases cell wall-deficient cells, called S-cells, under conditions of hyperosmotic stress ^32^. These S-cells are only transiently wall-deficient and can switch to the mycelial mode-of-growth. In some cases, however, prolonged exposure to high levels of osmolytes can lead to the emergence of mutants that are able to proliferate in the wall-deficient state as so-called L-forms ^32,33^. Like S-cells, these L-forms retain the ability to construct functional peptidoglycan based on the observation that removal of the osmolytes from the medium led to the formation of mycelial colonies. L-forms can also be generated in most other bacteria by exposing cells to compounds that target the process of cell wall synthesis ^33–35^. Strikingly, such wall-deficient cells that are able to propagate without the FtsZ-based cell division machinery ^35–37^. Even though the procedures used to generate L-forms can markedly differ, their mode-of-proliferation is conserved across species and largely based on biophysical principles. An imbalance in the cell surface area to volume ratio in cells that increase in size causes strong deformations of the cell membrane, followed by the release of progeny cells by blebbing, tubulation and vesiculation ^32,38^. Given that lipid vesicles without any content are able to proliferate in a similar manner to that observed for L-forms led to the hypothesis that this mode of proliferation may be comparable to that used by early life forms that existed before the cell wall had evolved ^39,40^.

Here, we exploited the unique properties of a *K. viridifaciens* L-form strain that readily switches between a wall-deficient and filamentous mode-of-growth to discover a novel MurG-like enzyme that is important for building the PG-based cell wall. Our data surprisingly show that *K. viridifaciens* produces wild-type peptidoglycan in the absence of *murG*, which was so far considered essential for Lipid II biosynthesis in all bacteria. The MurG activity is taken over by a novel paralogue called MurG2, which occurs widespread in filamentous actinobacteria, and able to substitute for the absence of MurG across different genera.

## RESULTS

### Morphological transitions of the shape-shifting strain *alpha*

We recently generated a *K. viridifaciens* L-form lineage by exposing the parental wild-type strain to high levels of penicillin and lysozyme. This strain, designated *alpha*, proliferates indefinitely in the cell wall-deficient state in media containing high levels of osmolytes ^32^ (Table S1). On solid LPMA medium, *alpha* forms green-pigmented viscous colonies, which exclusively contain L-form cells (Fig. 1A). In contrast, the parental strain forms compact and yellowish colonies composed of mycelia and S-cells on LPMA medium (Fig. 1B). Likewise, in liquid LPB medium *alpha* exclusively proliferates in the wall-deficient state, in a manner that is morphologically similar to that described for other L-forms ^35,41,42^; (Extended Data Video S1; Fig. 1C). Following strong deformations of the mother cell membrane (see panels of 56, 150, and 200 min in Fig. 1C), small progeny cells are released after approximately 300 min. The mother cell, from which the progeny was released (indicated with an asterisk) lysed after 580 min. Characterization using transmission electron microscopy (TEM) confirmed that *alpha* possessed no PG-based cell wall when grown on media containing high levels of osmolytes (Fig. 1D). Notably, when *alpha* is plated on MYM medium (lacking high levels of osmolytes) the strain can switch to the mycelial mode-of-growth (Fig. 1E). However, unlike the wild-type strain (Fig. 1F), the mycelial colonies of *alpha* fail to develop aerial hyphae and spores. Subsequent transfer of mycelia to LPMA plates stopped filamentous growth and reinitiated wall-deficient growth, during which L-form cells are extruded from stalled hyphal tips (Extended Data Video S2; Fig. 1G). Given the ability of these wall-deficient cells to proliferate, they eventually dominated the culture (not shown). Taken together, these results demonstrate that *alpha* can switch between a walled and wall-deficient state.

**Figure 1.**
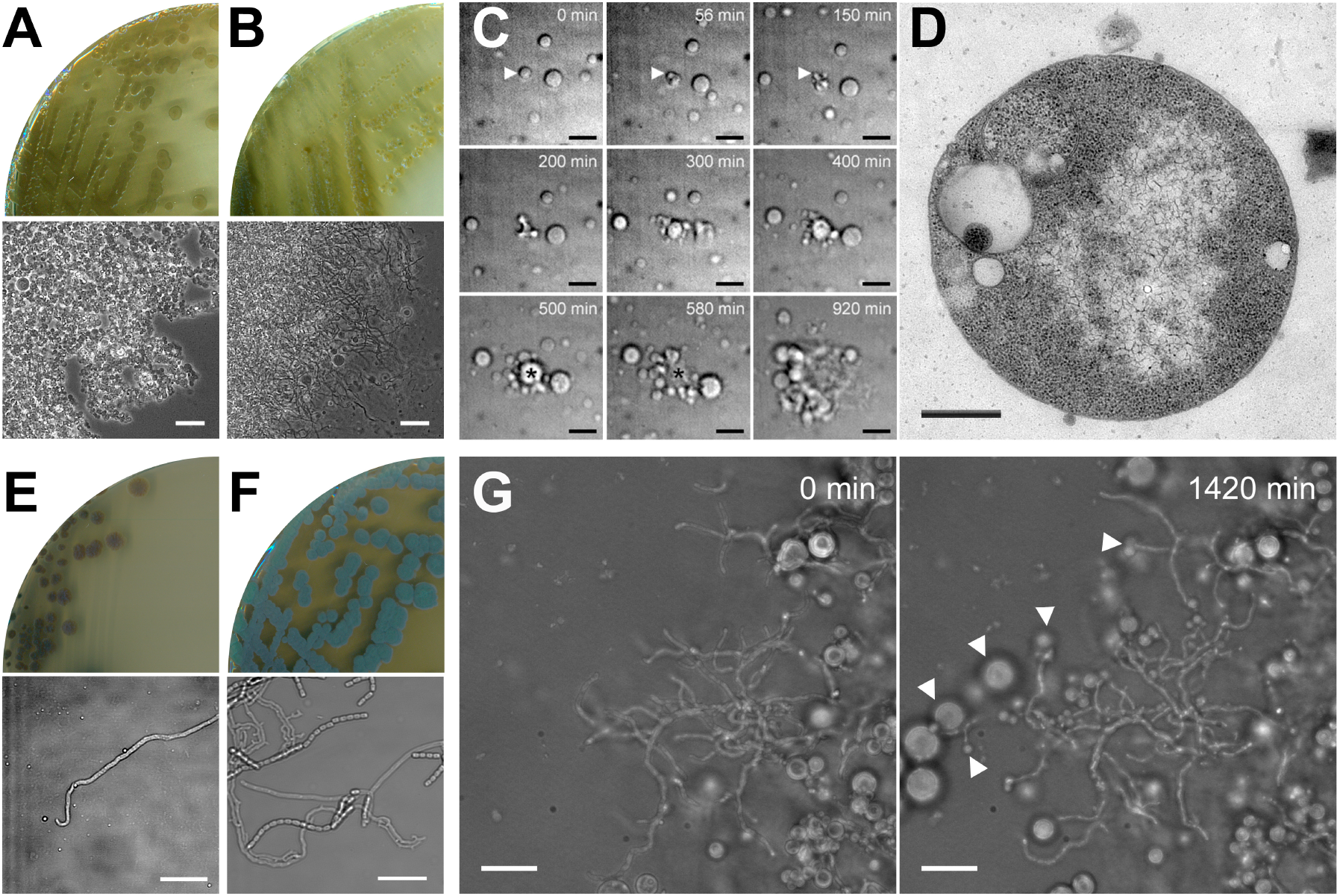
Morphological transitions of the shape-shifting strain *alpha*. (A) Growth of the *K. viridifaciens alpha* strain on LPMA medium yields green, mucoid colonies exclusively consisting of L-form cells, unlike the wild-type strain that forms yellowish colonies consisting of mycelia and S-cells (B). (C) Time-lapse microscopy stills of *alpha* proliferating in the wall-deficient state in liquid LPB medium. The arrowhead shows the mother cell, which generates progeny and lyses after 580 min (marked with an asterisk). Stills were taken from Supplementary Movie 1. (D) Transmission electron microscopy of a wall-deficient cell of *alpha*. (E) Growth of *alpha* on solid MYM medium yields compact, non-sporulating colonies unlike the wild-type strain that forms grey-pigmented sporulating colonies (F). (G) Time-lapse microscopy stills of mycelium of *alpha* transferred to LPMA medium, which show the extrusion of L-forms by filaments (see arrowheads). Stills were taken from Supplementary Movie 2. Scale bars represents 20 μm (A, B), 10 μm (C, E, F) and 500 nm (D).

### Deletion of *divIVA* abolishes switching of *alpha* from the wall-deficient to the filamentous mode-of-growth

The ability of *alpha* to efficiently switch between the walled and wall-deficient state provides an ideal platform to delete genes essential for cell wall biosynthesis. As a proof-of-concept, we focused on *divIVA*, which is essential for polar growth in filamentous actinomycetes ^24^. In Actinobacteria, *divIVA* is located adjacent to the conserved *dcw* gene cluster (Fig. S1). *divIVA* is present in Gram-positive rod-shaped (*Mycobacterium, Corynebacterium, Bacillus*), filamentous (*Streptomyces and Kitasatospora*) and coccoid (*Staphylococcus* and *Streptococcus*) bacteria, but absent in Gram-negatives such as *Escherichia coli*. In *B. subtilis* and *Staphylococcus aureus*, the DivIVA proteins share only 29% (BSU15420) and 26% (SAOUHSC_01158) aa identity to the *S. coelicolor* orthologue. To localize DivIVA, plasmid pKR2 was created, allowing constitutive expression of DivIVA-eGFP (Table S2). Fluorescence microscopy revealed that the fusion protein localized to hyphal tips (Fig. S2A), similarly as in streptomycetes ^24^. When *alpha* was grown in the wall-deficient state in LPB medium, typically one or two foci of DivIVA-eGFP were detected per cell, which invariably were localized to the membrane. In contrast, no foci were detected in L-form cells containing the empty plasmid (pKR1) or those expressing cytosolic eGFP (pGreen ^43^). We then constructed the plasmids pKR3 to delete *divIVA* and pKR4 to delete a large part of the *dcw* gene cluster, including *divIVA* (Table S2). Introduction of these plasmids into *alpha* by PEG-mediated transformation and a subsequent screening yielded the desired *divIVA* and *dcw* mutants (Fig. S3). Analysis of growth in LPB medium or on solid LPMA plates indicated that the L-form cells proliferated normally in the absence of *divIVA* or part of the *dcw* gene cluster (Fig. 2). However, when L-form cells were plated on MYM medium (lacking osmoprotectants), only the *alpha* strain was able to switch to the mycelial mode-of-growth (Fig. 2B). Introduction of plasmid pKR6, which expresses *divIVA* from the constitutive *gap1* promoter, complemented growth of the *divIVA* mutant on MYM medium (Fig. 2B). In agreement, Western blot analysis using antibodies against DivIVA of *Corynebacterium glutamicum* confirmed the absence of DivIVA in both the *divIVA* and the *dcw* mutant, and also showed the expression was restored in the *divIVA* mutant complemented with pKR6 (Fig. 2C).

**Figure 2.**
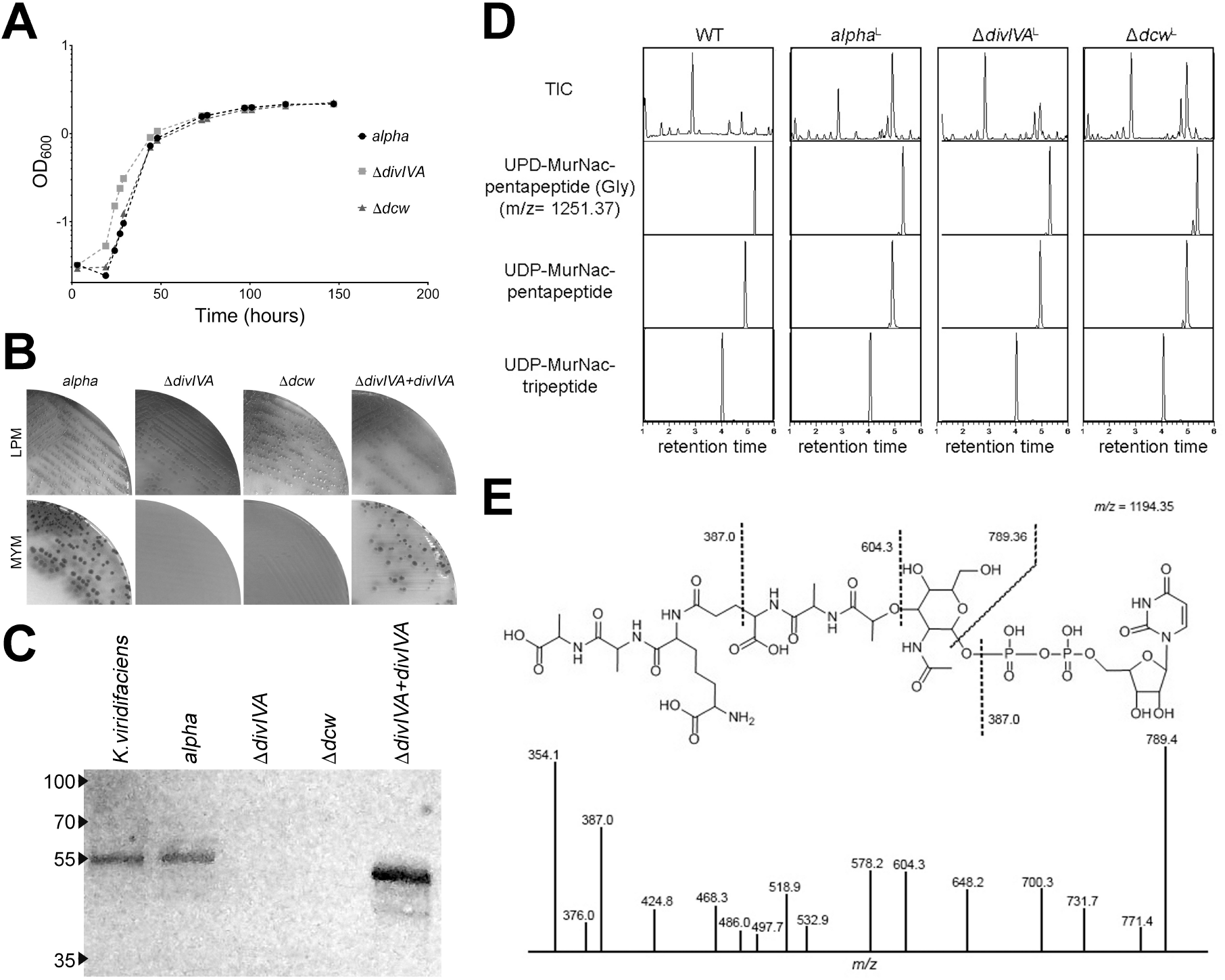
The absence of DivIVA abolishes switching of *alpha* from the wall-deficient to the filamentous mode-of-growth. (A) Growth curves of *alpha* (black spheres), the *divIVA* mutant (grey squares) and the *dcw* mutant (grey triangles) in liquid LPB medium. (B) While all strains grow on LPMA medium, those lacking *divIVA* are unable to switch to the mycelial mode-of-growth on MYM medium lacking osmoprotectants. (C) Western Blot analysis using antibodies against the *C. glutamicum* DivIVA protein confirm the absence of DivIVA in the constructed *ΔdivIVA* and *dcw* mutants. Reintroduction of *divIVA* under control of the *gap1* promoter restores the expression of DivIVA in the *divIVA* mutant and the ability to form mycelial colonies (see panel B). (D) Comparative LC-MS analysis of peptidoglycan precursors in *alpha* and its *divIVA* and *dcw* mutants. Like the wild-type, all strains produce peptidoglycan precursors including UDP-MurNAc-pentapeptide, which is the last cytosolic precursor in the PG biosynthesis pathway. (E) MS-MS analysis demonstrating that the product with a mass of 1194.35 is the precursor UDP-MurNAc-pentapeptide.

To analyse if the switch from the wall-deficient to the walled state in the absence of DivIVA was blocked due to the failure to produce the cytosolic precursors required for peptidoglycan synthesis in the L-form state, we performed a comparative LC-MS analysis (Fig. 2D). We noticed that the LC-MS profiles of the *divIVA* and *dcw* mutant strains were similar to that of *alpha* with respect to the cytosolic PG building blocks (Fig. 2D). Importantly, MS-MS analysis identified the last cytosolic precursor in the PG biosynthesis pathway, UDP-MurNAc-pentapeptide (Mw = 1194.35) in all strains (Fig. 2E). Taken together, these results demonstrate that DivIVA is essential for filamentous growth but not required for synthesis of the cytosolic PG precursors.

### Identification of a distant MurG paralogue as a novel Lipid II synthase

Having a mutant lacking many genes of the *dcw* cluster offers many opportunities for the study of individual genes. The constructed *dcw* mutant lacks *ftsW, murG, ftsQ, ftsZ, ylmD, ylmE, sepF, sepG*, and *divIVA*. Surprisingly, introduction of only *divIVA* (expressed from the constitutive *gap1* promoter) restored the ability of the *dcw* mutant to switch to the walled mode-of-growth on solid media lacking osmoprotectants (Fig. 3). The colonies that were formed, were small and heterogeneous as compared to the mycelial colonies formed by *alpha* (Fig. 3A). Furthermore, expression of *divIVA* in the *dcw* mutant was not able to restore filamentous growth in liquid cultures (data not shown). To verify that the *dcw* mutant expressing *divIVA* produced normal PG on solid medium, we performed a peptidoglycan architecture analysis using LC-MS (Fig. 3B). This surprisingly revealed that all expected muropeptides were formed at levels comparable to those formed by *alpha* and the wild-type strain, despite the absence of a functional *murG* (Fig. 3B; Table 1).

**Figure 3.**
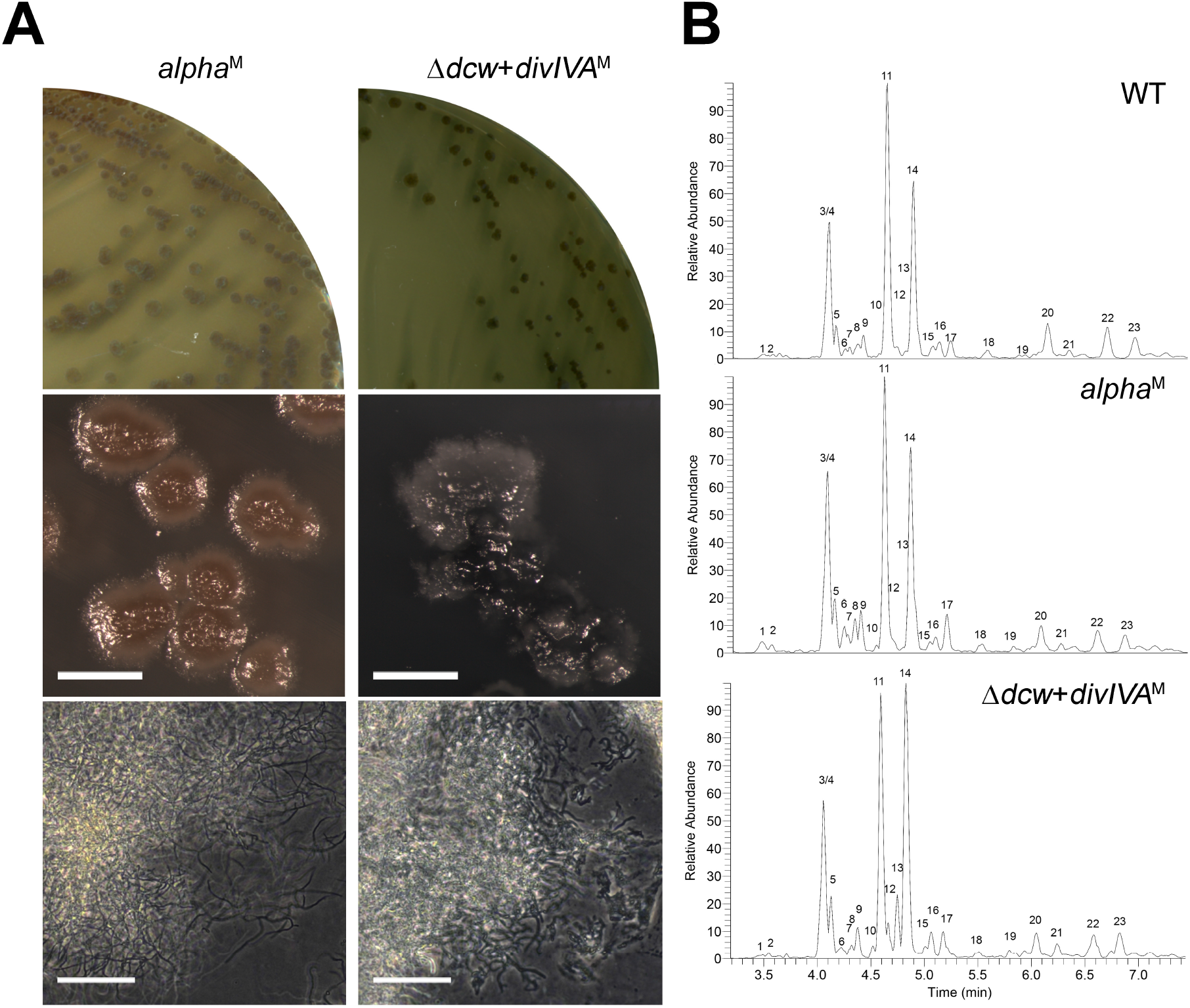
Reintroduction of *divIVA* alone is sufficient to restore filamentous growth of the *dcw* mutant. (A) Morphological comparison between *alpha* (left) and the *dcw* mutant transformed with *P_gap1_-divIVA* (right) grown on MYM medium. Unlike *alpha*, the *dcw* mutant expressing DivIVA forms colonies with a heterogenous appearance. (B) Peptidoglycan architecture analysis of mycelium of the wild-type strain (top), *alpha* (middle) and the *dcw* mutant expressing DivIVA (bottom). The abundance of muropeptides is similar in all strains despite the lack of *murG* in the *dcw* mutant (see also Table 1). Scale bar, 40 μm.

**Table 1.**
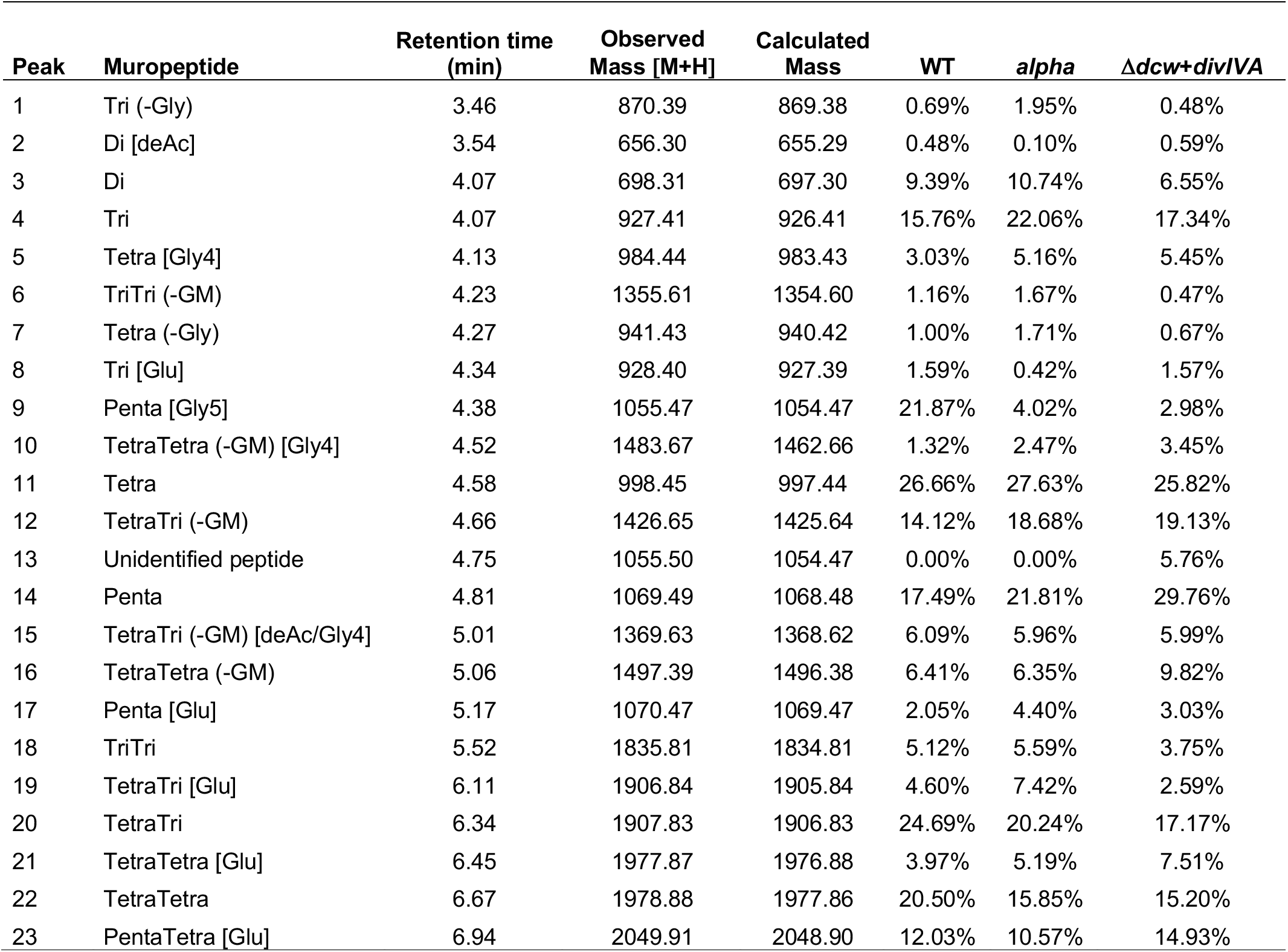
Muropeptides identified in *K. viridifaciens* strains grown as mycelium. Monomers and dimers are treated as separate sets. Masses are indicated in Da.

The ability of the *dcw* mutant expressing *divIVA* to grow filamentous inevitably means that another protein had functionally replaced the activity of MurG. Blast analysis of the amino acid sequence of MurG_SCO_ (SCO2084) against the genome sequence of *K. viridifaciens* revealed that this actinomycete contains two putative, but distant MurG homologs (Supplemefntary Table 4). The two additional homologs (BOQ63_RS12640 and BOQ63_RS05415) showed 31.2% and 16.5% sequence identity, respectively, to MurG (BOQ63_RS32465). Further investigation revealed that MurG proteins possess two characteristic domains: an N-terminal domain that contains the Lipid I binding site (PF03033) ^44^, and a C-terminal domain that contains the UDP-GlcNAc binding site (PF04101; Fig. S4), both of which are required for the UDP-N-acetyl-glucosamine transferase activity. Of the two distant MurG homologues, only BOQ63_RS12640 contained both domains (Fig. S4). A broader search of MurG-like proteins in other *Streptomyces* and *Kitasatospora* spp. revealed that 38% of the strains possess one, two and sometimes even three genes for MurG-like proteins containing both the necessary N-terminal (PF03033) and C-terminal (PF04101) domains (Fig. 4), in addition to the canonical MurG, which is present in all strains and encoded in the *dcw* gene cluster. A sequence similarity network was produced by pairwise comparing the 1553 MurG and MurG-like proteins extracted from all translated *Streptomyces* and *Kitasatospora* genomes, which showed that nearly all MurG proteins encoded by the orthologue of *murG* in the *dcw* gene cluster grouped together. However, the MurG-like proteins clustered in many different groups (Fig. S5).

**Figure 4.**
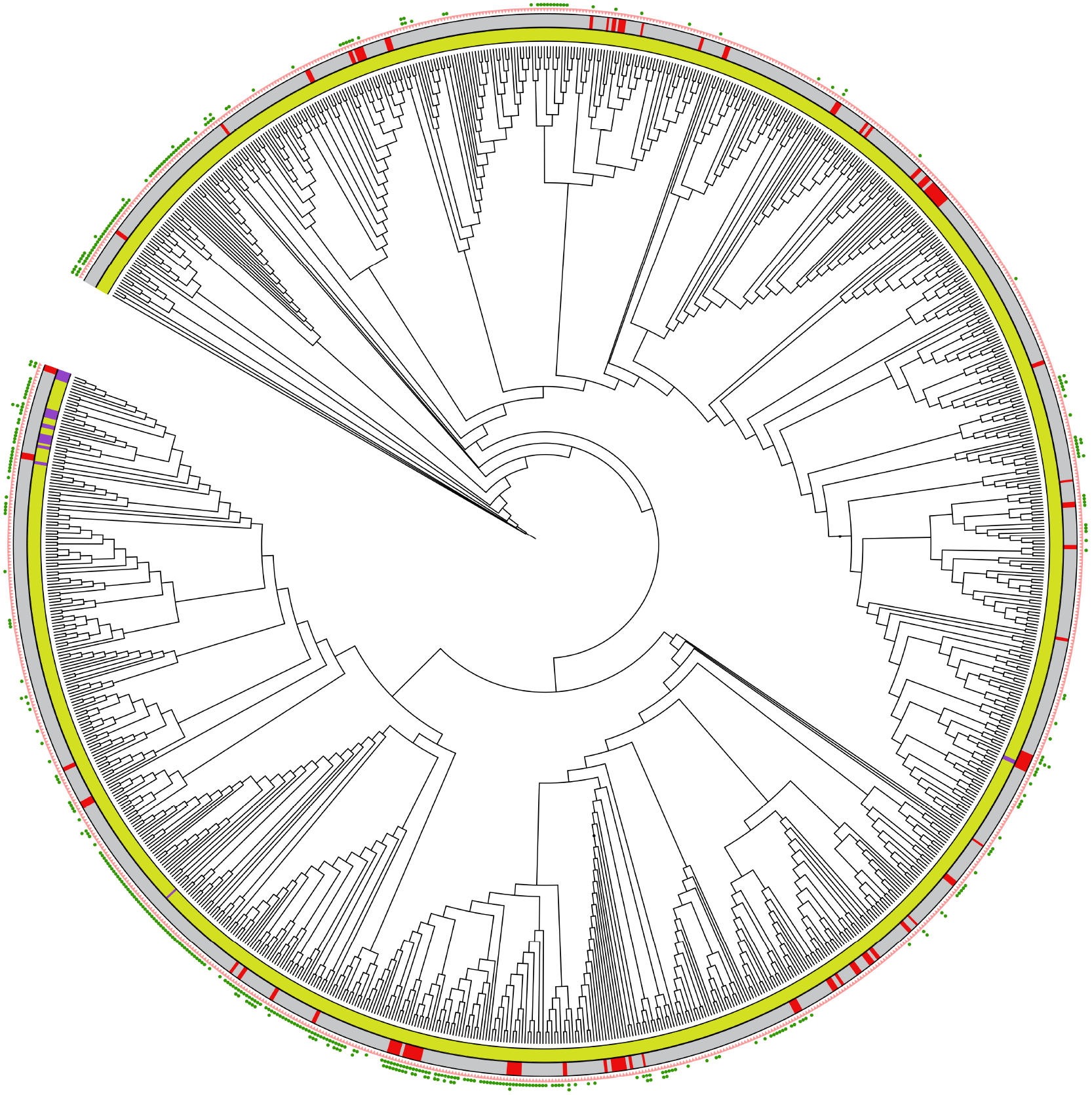
Overview of MurG and MurG-like proteins present in *Streptomyces* and *Kitasatospora* species. The phylogenetic tree was constructed on the basis of four 4 conserved housekeeping proteins (AtpD, RecA, TrpB and GyrB). Yellow and purple colors in the inner circle represent *Streptomyces* and *Kitasatospora* species, respectively. Strains present in the NCBI database are indicated in grey in the middle circle, while those from an in-house collection are indicated in red. The pink triangles represent MurG proteins encoded in the *dcw* gene cluster. The green dots represent distant MurG proteins, whose genes are located elsewhere in the genomes. Phylogenetic trees were constructed using iTOL ^70^.

To corroborate that *murG* is not required for filamentous growth, we decided to delete *murG* in *alpha* using knock-out construct pKR8 (Table S2). The genotype of the mutant was verified by PCR (Fig. S6) and showed that the absence of *murG* had no effect on L-form or filamentous growth (Fig. 5). Likewise, inactivation of *murG2* in *alpha* using construct pKR9 had no effect on L-form growth and did not prevent switching to mycelial growth (Fig. 5). We then attempted to create a double mutant by deleting *murG2* in the *murG* mutant. PCR analysis on a putative double mutant strain with the highly sensitive Q5 DNA polymerase indicated, however, that a small proportion of the multinucleated L-forms had retained a copy of *murG2* (Fig. S6). Also, further subculturing of this merodiploid strain in the presence of antibiotics that counter-selected for maintenance of *murG2* did not lead to a complete loss of this gene, suggesting that the ability to produce Lipid II is essential in these L-forms (see Discussion). Nevertheless, plating this merodiploid strain on MYM medium essentially blocked mycelial growth, and only at very high cell densities infrequent shifters were found (see encircled colony in Fig. 5A).

**Figure 5.**
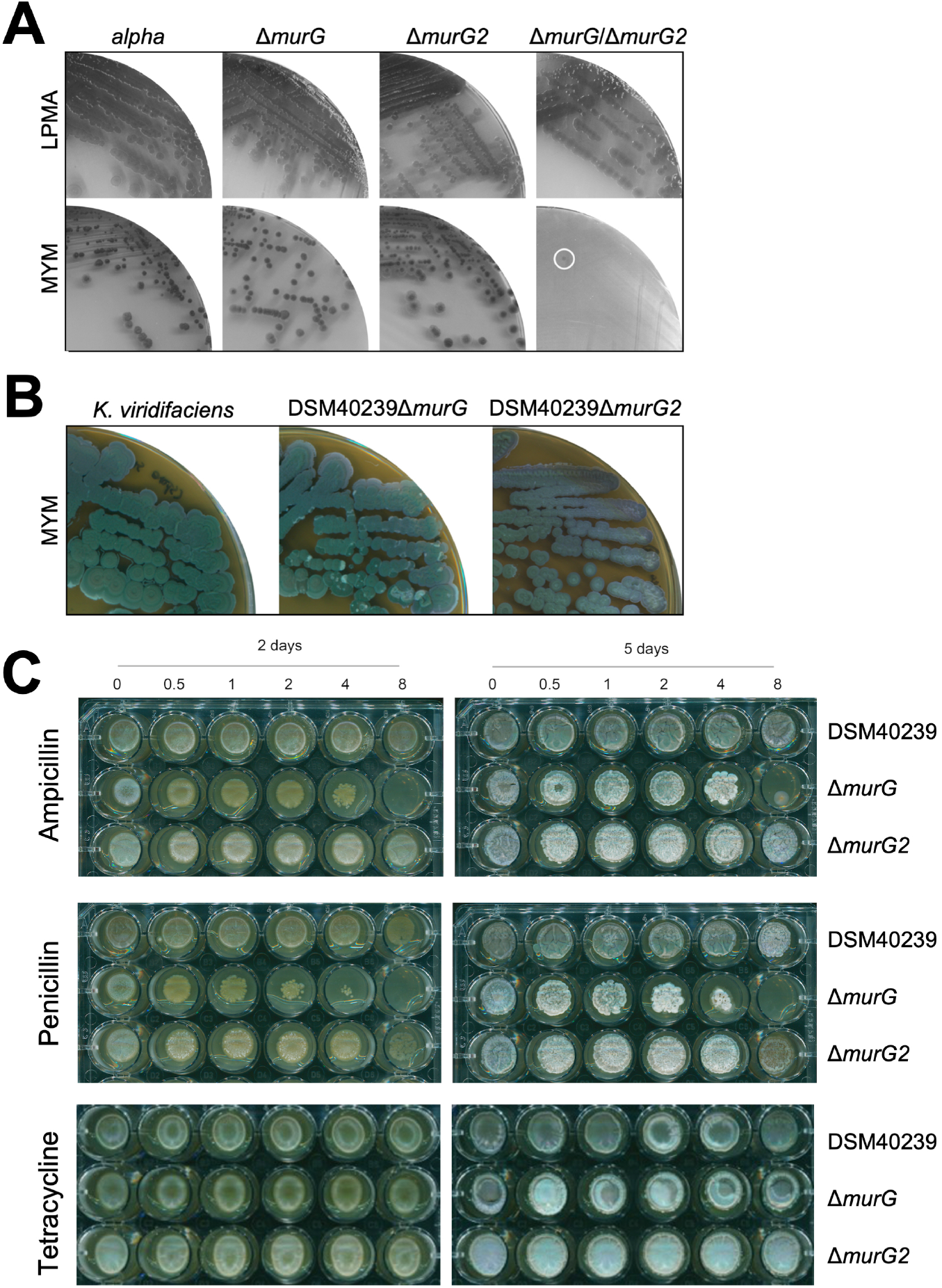
MurG2 can functionally replace MurG in peptidoglycan synthesis. (A) Plates of *alpha* and the Δ*murG*, Δ*murG2* and the merodiploid Δ*murG*Δ*murG2* strains on LPMA medium (top). With the exception of the Δ*murG*Δ*murG2* merodiploid, all strains efficiently switched to filamentous growth on MYM medium lacking osmolytes (bottom). (B) Plates of *K. viridifaciens* and its Δ*murG* and Δ*murG2* mutants grown on MYM medium for 7 days. (C) Plates of *K. viridifaciens* and the Δ*murG* and Δ*murG2* mutant strains grown on MYM medium for 2 (left) or 5 (right) days in the presence of ampicillin (top), penicillin (middle) and tetracycline (bottom). The antibiotic concentrations (in μg ml^−1^) are indicated above the plates.

Having demonstrated that *murG* is not required for filamentous growth of *alpha*, we then wondered whether *murG* would also be dispensable for filamentous growth of the wild-type strain. Notably, *murG* deletion mutants could not be obtained if transformants were selected on MYM medium, unlike a *murG2* deletion mutant that was readily found. However, when transformants were selected on LPMA medium containing high levels of sucrose, a *murG* mutant could be created in *K. viridifaciens* (see Fig. S7). As shown in Figure 5B, the generated *murG* and *murG2* mutants were able to develop and sporulate normally on MYM, when compared to the parental wild type. However, exposing the strains to low levels of penicillin and ampicillin revealed that the *murG* mutant was more susceptible to these cell wall-targeting antibiotics when compared to the wild-type and its *murG2* mutant. By contrast, no difference effect was observed when tetracycline was added to the plates (Fig. 5C). Altogether, these results demonstrate that MurG and MurG2 have overlapping activities, whereby MurG2 is able to functionally replace the canonical Lipid II synthase MurG.

### MurG2 from *K. viridifaciens* can functionally replace MurG in *S. coelicolor*

The observations that *murG2* can functionally replace *murG* in *K. viridifaciens* and that strains expressing only MurG2 produce wild-type peptidoglycan, strongly suggest that the *murG2* gene product synthesizes Lipid II. To further substantiate this, we investigated whether *murG2* could also functionally complement *murG* (SCO2084) in another Actinobacterium, namely the model organism *Streptomyces coelicolor* M145, which itself does not harbour an orthologue of *murG2*. For this, we created construct pGWS1379 expressing *murG2* from the constitutive *ermE* promoter in the integrative vector pMS82 and introduced it into *S. coelicolor*. As a control we used the empty vector pMS82. We then applied CRISPRi ^45^ to knock-down the native *murG* to assess viability. CRISPRi only works when the spacer of the dCas9/sgRNA complex targets the non-template strand of *murGsco*, and not the template strand, or when the spacer is absent ^45,46^. The functionality of the CRISPRi constructs was evident in control cells without *murG2;* colonies expressing the dCas9/sgRNA complex targeting the non-template strand of *murGsc* in M145 could hardly grow, due to the essential function of *murG*. Conversely, control transformants harboring CRISPRi constructs targeting the template strand or without spacer (empty plasmid) grew normally (Fig. 6). Excitingly, *S. coelicolor* transformants expressing *murG2* grew apparently normally under all conditions, even when *murG* expression was knocked down by the CRISPRi system. This validates the concept that *murG2* can functionally replace the canonical *murG* (Fig. 6). Taken together, our experiments show that the MurG2 enzyme can functionally replace the Lipid II-biosynthetic enzyme MurG, both in *Kitasatospora* and in *Streptomyces*.

**Figure 6.**
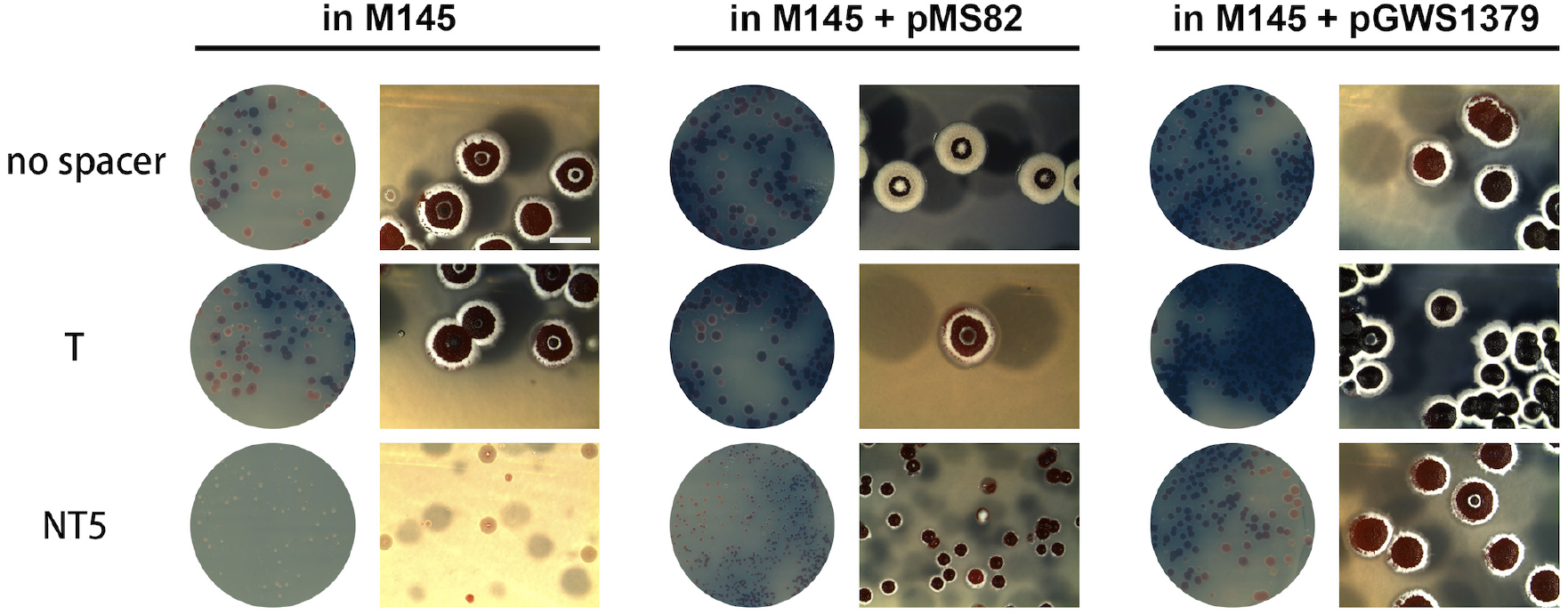
Ectopic expression of *murG2* allows silencing of *murGsc* via CRISPRi. CRISPRi constructs were introduced into *S. coelicolor* M145 or with control plasmid pMS82 and a recombinant strain with pGWS1372 integrated in its genome, thus expressing *K. viridifaciens* MurG2. Expectedly, no effect was seen when CRISPRi constructs were introduced that either had no spacer or that contained a spacer targeting the template strand (T) of *murGsc*. However, constructs targeting the non-template strand (NT) resulted in severe phenotypic defects and sick colonies of *S. coelicolor* that lacked *murG2*, but not in pGWS1379 transformants that expressed *murG2*. Images were taken after 5 days incubation at 30°C. Bar, 2 mm.

## DISCUSSION

The cell wall is a hallmark feature of bacterial cells, and the steps involved in its biosynthesis are widely conserved across the bacterial domain. In all bacteria, the final cytosolic step in precursor biosynthesis is the conversion of Lipid I to Lipid II by MurG encoded in the *dcw* gene cluster. We here show for the first time that the novel enzyme MurG2 can replace the activity of MurG and demonstrate that *murG* is dispensable in the filamentous actinomycete *K. viridifaciens* in the presence of *murG2*. MurG2 alone is sufficient to produce wild-type peptidoglycan. MurG2 is in fact widespread among the *Streptomycetaceae* and was identified in the genomes of 38% of all *Streptomyces* and *Kitasatospora* strains. Furthermore, introduction of *K. viridifaciens murG2* into *S. coelicolor* M145 – which itself lacks an orthologue of *murG2* – allowed the knock-down of the canonical *murG* using CRISPRi, showing that the gene is a *bona fide* cell wall biosynthetic gene that is functional in different Actinobacteria.

Filamentous actinomycetes are multicellular bacteria that form networks of interconnected hyphae, whereby sporulating aerial hyphae are established after a period of vegetative growth. *Streptomyces* is a wonderful model system for the study of cell division, among others because cell division is not required for normal growth of this bacterium ^21,25,47^. Most of the cell division proteins are encoded by genes located in the conserved *dcw* gene cluster. In streptomycetes, many cell division genes such as *ftsI, ftsL, ftsW* and *divIC* are only required for sporulation and do not affect normal growth ^48–50^. Our data surprisingly show that many genes within the *dcw* cluster can be deleted simultaneously in *K. viridifaciens*, including *divIVA* that is essential for polar growth in Actinobacteria, by using a strain (*alpha*) with the ability to readily switch between a wall-deficient and filamentous mode-of-growth. The *alpha* strain thus provides a unique system for the identification of proteins that are required for polar growth. As a proof-of-concept for this principle, *divIVA* that is required for polar growth, was successfully deleted. Consistent with its role in driving apical growth, the absence of *divIVA* arrested growth in the wall-deficient state but had no effect on synthesis of the PG building blocks. This indicates that the block in PG formation occurred in a later step of the PG biosynthesis pathway. Introduction of only *divIVA* in the *dcw* mutant retored polar growth, which was a rather surprising discovery given the absence of a whole string of genes involved in cell division and cell wall synthesis, and in particular *murG*. MurG catalyzes the coupling of GlcNAc to Lipid I, yielding the PG precursor Lipid II and this enzymatic activity is therefore essential for cell wall synthesis. The ability of *alpha* to produce a cell wall with an apparently normal architecture, as shown by the analysis of the peptidoglycan, indicated that *K. viridifaciens* possesses other enzymes capable of synthesizing Lipid II in the absence of *murG*. An in-silico search in the genome of *K. viridifaciens* identified *murG2* (BOQ63_RS12640), which is a distant relative of MurG with the likely ability to replace the activity of the canonical MurG. This is based among others on the presence of the two domains that are known to be required for the transfer of GlcNAc to Lipid I. Many Actinobacteria possess proteins carrying these two domains, suggesting that MurG2 proteins are common in these bacteria. In fact, some species even contain three genes for MurG-like proteins, in addition to the canonical MurG encoded in the *dcw* gene cluster. Interestingly, *murG* and *murG2* could both be individually deleted in the wild-type strain, whereby the resulting mutants showed normal growth and development when strains were grown in non-stressed environments. However, the *murG* mutant was more susceptible to cell wall-targeting antibiotics than the wild-type strain or its *murG2* mutant. Considering that MurG2 alone suffices to produce normal peptidoglycan, this suggests that MurG is required to build a more robust cell wall. Deletion of *murG* was only possible after exposing transformants to hyperosmotic growth conditions. We hypothesize that the hyperosmotic conditions activated the transcription of *murG2*, thus allowing deletion of *murG* specifically under these growth conditions. This implies that the function of *murG2* is to synthesize Lipid II under specific growth conditions, e.g. during hyperosmotic stress.

In further support of the function of MurG2 as an alternative Lipid II synthase, we tested if it could also take over the function of *murG* in another bacterium. For this, we chose the model streptomycete *S. coelicolor* M145, which is a distinct genus within the *Streptomycetaceae* ^31,51^, but lacking a copy of *murG2*. Importantly, *murG* could be readily depleted using CRISPRi in strains expressing *murG2* from a constitutive promoter, while knock-down of *murG* in colonies of *S. coelicolor* harboring control plasmids led to very severe growth defects. This not only validates our data that *murG2* encodes a Lipid II synthase, but also that this is a more universal phenomenon that does not only occur in specific strains of *Kitasatospora* or connect to strains that have the capacity to produce natural wall-less cells. Furthermore, it shows that no additional *Kitasatospora* genes are required to allow *murG2* to functionally complement *murG* in *Streptomyces*.

We also attempted to delete *murG* and *murG2* simultaneously in *alpha*. While the single mutants were readily obtained, we never obtained strains that completely devoid of both *murG* and *murG2*, despite many attempts. Like mycelia, L-forms are multinucleated cells, and some cells of the population retained *murG2*, most likely to ensure minimal levels of Lipid II. Consistent with this idea is the finding that antibiotics that target Lipid II, such as vancomycin, are lethal to *alpha* (our unpublished data). We hypothesize that this lethality is caused by depletion of the lipid carrier undecaprenyl diphosphate, which is also used in other pathways and which may be essential for these L-forms. Removing *murG2* in strains lacking *murG* strain virtually blocked the ability to switch to the filamentous mode-of-growth, whereas each of the single mutants switched as efficiently as the parental *alpha* strain. Thus, we show that MurG2 is a novel enzyme involved in cell wall metabolism, which appears to facilitate switching between a wall-deficient and a walled lifestyle.

## MATERIALS AND METHODS

### Strains and media

Bacterial strains used in this study are shown in Table S1. To obtain sporulating cultures of *K. viridifaciens* and *S. coelicolor*, strains were grown at 30°C for 4 days on MYM medium ^52^. For general cloning purposes, *E. coli* strains DH5α and JM109 were used, while *E. coli* ET12567 and SCS110 were used to obtain unmethylated DNA. *E. coli* strains were grown at 37 °C in LB medium, supplemented with chloramphenicol (25 μg ml^−1^), ampicillin (100 μg ml^−1^), apramycin (50 μg ml^−1^), kanamycin (50 μg ml^−1^), or viomycin (30 μg ml^−1^), where necessary.

To support growth of wall-deficient cells, strains were grown in liquid LPB medium while shaking at 100 rpm, or on solid LPMA medium at 30°C ^32^. To switch from the wall-deficient to the filamentous mode-of-growth, L-form colonies grown on LPMA for seven days were streaked on MYM medium. If needed, mycelial colonies of switched strains were transferred after 4 days to liquid TSBS medium and grown for two days at 30°C, while shaking at 200 rpm.

### Construction of plasmids

All plasmids and primers used in this work are shown in Tables S2 and S3, respectively.

#### Construction of the DivIVA localization construct pKR2

To localize DivIVA, we first created plasmid pKR1 containing a viomycin resistance cassette cloned into the unique NheI site of pIJ8630 ^53^. To this end, the viomycin resistance cassette was amplified from pIJ780 ^54^ with the primers *vph*-FW-NheI and *vph*-RV-NheI. Next, we amplified the constitutive *gap1* promoter as a 450 bp fragment from the genome of *S. coelicolor* with the primers P*gap1*-FW-BglII and P*gap1*-RV-XbaI. We also amplified the *divIVA* coding sequence (the +1 to +1335 region relative to the start codon of *divIVA* (BOQ63_RS32500) from the chromosome of *K. viridifaciens* using primers *divIVA*-FW-XbaI and *divIVA*-Nostop-RV-NdeI ^55^. Finally, the promoter and *divIVA* coding sequence were cloned into pKR1 as a BglII/XbaI and XbaI/NdeI fragment respectively, yielding plasmid pKR2.

#### Construction of the deletion constructs pKR3, pKR4, pKR8, pKR9 and pKR10

The *divIVA* mutant was created in *K. viridifaciens* using pKR3, which is a derivative of the unstable plasmid pWHM3 ^56^. In the *divIVA* mutant, nucleotides +205 to +349 relative to the start codon of *diviVA* were replaced with the *loxP-apra* resistance cassette as described ^57^. A similar strategy was used for the deletion of the partial *dcw* cluster (plasmid pKR4), and for the deletion of *murG* (plasmid pKR8) and *murG2* (plasmid pKR9). For the deletion of the partial *dcw* cluster, the chromosomal region from +487 bp relative to the start of the *ftsW* gene (BOQ63_RS32460) until +349 relative to the start of the *divIVA* gene were replaced with the apramycin resistance marker. For the deletion of *murG* (BOQ63_RS32465, located in the *dcw* cluster), the nucleotides +10 to +1077 bp relative to the start codon of *murG* were replaced with the *loxP-apra* resistance cassette, while for the *murG2* (BOQ63_RS12640) deletion the chromosomal region from +18 to +1105 bp relative to the start of *murG2* were replaced with the apramycin resistance marker. To construct the *murG/murG2* double mutant, pKR10 was created, replacing the apramycin resistance cassette in pKR8 by a viomycin resistance cassette. To this end, the viomycin resistance cassette was amplified from pIJ780 ^54^ with the primers *vph*-Fw-EcoRI-HindIII-XbaI and *vph*-Rv-EcoRI-HindIII-XbaI. The viomycin resistance cassette contained on the PCR fragment was then cloned into pKR8 using XbaI, thereby replacing the apramycin cassette and yielding pKR10.

#### Construction of the complementation constructs pKR6 and pKR7

For complementation of *divIVA* under control of the strong *gap1* promoter ^43^, the constructs pKR6 was made. First, we created plasmid pKR5 with the strong *gap1* promoter. The promoter region of *gap1* (SCO1947) was amplified with the primers P*gap1*-FW-BglII and P*gap1*-RV-XbaI using *S. coelicolor* genomic DNA as the template. Next, the *gap1* promoter was cloned as BglII/XbaI fragment into the integrative vector pIJ8600 ^53^ to generate the plasmid pKR5. Afterwards, the *divIVA* coding sequence was amplified from the genome of *K. viridifaciens* with the primers *divIVA*-XbaI-FW and *divIVA*-NdeI-RV. Finally, to create the plasmid pKR6 the XbaI/NdeI fragment containing the *divIVA* coding sequence was cloned in pKR5.

#### Construction of the murG2 expression construct pGWS1379

A DNA fragment containing the *ermE\** promoter was obtained as an EcoRI-NdeI fragment from pHM10a ^58^, while *murG2* was amplified by PCR from *K. viridifaciens* chromosomal DNA using primer pair murG2_F+4_ENdeI and murG2_R+1146_HX. The *ermEp\** fragment and NdeI-XbaI-digested *murG2* were simultaneously cloned into EcoRI-XbaI digested pSET152 to generate construct pGWS1378. The insert of pGWS1378 was then introduced as a PvuII fragment into EcoRV-digested pMS82 ^59^ to generate construct pGWS1379. This construct was then introduced into *S. coelicolor* M145 via protoplast transformation as described ^60^.

### Transformation of L-forms

Transformation of *alpha* essentially followed the protocol for the rapid small-scale transformation of *Streptomyces* protoplasts ^60^, with the difference that 50 μl cells from a mid-exponential growing L-form culture were used instead of protoplasts. Typically, 1 μg DNA was used for each transformation. Transformants were selected by applying an overlay containing the required antibiotics in P-buffer after 20 hours. Further selection of transformants was done on LPMA medium supplemented with apramycin (50 μg ml^−1^), thiostrepton (5 μg ml^−1^), or viomycin (30 μg ml^−1^), when necessary. Transformants were verified by PCR (Table S3).

### *murGsco* (SCO2084) knockdown via CRISPRi

The NcoI restriction site within the integrase gene of phage φC31 in pSET152 was removed by introducing a silent GCC to GCG change in codon A360 via site-directed mutagenesis by PCR using primer pairs 152DNcoI_F and 152DNcoI_R, to generate construct pGWS1369. Subsequently, a DNA fragment containing the sgRNA scaffold (no spacer) and *Pgapdh-dcas9* of constructs pGWS1049 ^46^ was cloned as an EcoRI-XbaI fragment into pGWS1369 to generate construct pGWS1370. The 20 nt spacer sequence was introduced into sgRNA scaffold by PCR using forward primers SCO2084_T_F or SCO2084_NT5_F together with the reverse primer SgTermi_R_B. The PCR products were cloned as NcoI-BamHI fragments into pGWS1370 to generate constructs pGWS1371 (targeting the template strand of SCO2084) and pGWS1376 (targeting the non-template strand of SCO2084). Constructs pGWS1370 (no spacer), pGWS1371 (targeting the template strand) and pGWS1376 (targeting the non-template strand) were introduced into *S. coelicolor* M145+pMS82 (empty plasmid) and M145+pGWS1379 (expressing *murG2*) via protoplast transformation as described previously ^60^.

### Microscopy

Strains grown in LPB or LPMA were imaged using a Zeiss Axio Lab A1 upright microscope equipped with an Axiocam Mrc. A thin layer of LPMA (without horse serum) was applied to the glass slides to immobilize the cells prior to the microscopic analysis.

#### Fluorescence microscopy

Fluorescence microscopy pictures were obtained with a Zeiss Axioscope A1 upright fluorescence microscope equipped with an Axiocam Mrc5 camera. Aliquots of 10 μl of live cells were immobilized on top of a thin layer of LPMA (without horse serum) prior to analysis. Fluorescent images were obtained using a 470/40 nm band pass excitation and a 505/560 band pass detection, using an 100x N.A. 1.3 objective. To obtain a sufficiently dark background, the background of the images was set to black. These corrections were made using Adobe Photoshop CS5.

#### Time-lapse microscopy

To visualize the proliferation of *alpha*, cells were collected and resuspended in 300 μl LPB (containing 4-22% sucrose) and placed in the wells of a chambered 8-well μ-slide (ibidi®). Cells were imaged on a Nikon Eclipse Ti-E inverted microscope equipped with a confocal spinning disk unit (CSU-X1) operated at 10,000 rpm (Yokogawa), using a 40x Plan Fluor Lens (Nikon) and illuminated in bright-field. Images were captured every 2 minutes for 10-15 hours by an Andor iXon Ultra 897 High Speed EM-CCD camera (Andor Technology). Z-stacks were acquired at 0.2-0.5 μm intervals using a NI-DAQ controlled Piezo element. During imaging wall-less cells were kept at 30 °C using an INUG2E-TIZ stage top incubator (Tokai Hit).

#### Electron microscopy

For transmission electron microscopy, L-forms obtained from a 7-day-old liquid-grown *alpha* culture were trapped in agarose blocks prior to fixation with 1.5% glutaraldehyde and a post-fixation step with 1% OsO4. Samples were embedded in Epon and sectioned into 70 nm slices. Samples were stained using uranyl-acetate (2%) and lead-citrate (0.4%), if necessary, before being imaged using a Jeol 1010 or a Fei 12 BioTwin transmission electron microscope.

### DivIVA detection using Western analysis

To detect DivIVA using Western analysis, the biomass of L-form strains was harvested after 7 days of growth in LPB medium, while biomass of mycelial strains was obtained from liquid-grown TSBS cultures after 17 hours. Cell pellets were washed twice with 10% PBS, after which they were resuspended in 50 mM HEPES pH 7.4, 50 mM NaCl, 0.5% Triton X-100, 1 mM PFMS and P8465 protease inhibitor cocktail (Sigma). The cells and mycelia were disrupted with a Bioruptor Plus Sonication Device (Diagenode). Complete lysis was verified by microscopy, after which the soluble cell lysate was separated from the insoluble debris by centrifugation at 13,000 rpm for 10 min at 4°C. The total protein concentration in the cell lysates was quantified by a BCA assay (Sigma-Aldrich). Equal amounts of total proteins were separated with SDS-PAGE using 12,5% gels. Proteins were transferred to polyvinylidene difluoride (PVDF) membranes (GE Healthcare) with the Mini Trans-Blot® Cell (Bio-Rad Laboratories) according to the manufacturer’s instructions. DivIVA was detected using a 1:5,000 dilution of polyclonal antibodies raised against *Corynebacterium glutamicum* DivIVA (kindly provided by Professor Marc Bramkamp). The secondary antibody, anti-rabbit IgG conjugated to alkaline phosphatase (Sigma), was visualized with the BCIP/NBT Color Development Substrate (Promega).

### Isolation of cytoplasmic peptidoglycan precursors

For the cytoplasmic PG precursor isolation and identification, we used a modification of the method previously described ^61^. The *alpha* strain and the *divIVA* and *dcw* mutants were grown in LPB for seven days, while the wild-type *K. viridifaciens* strain was grown for three days in a modified version of LPB lacking sucrose. The cells were harvested by centrifugation at 4°C and washed in 0,9% NaCl. Cells were extracted with 5% cold trichloric acid (TCA) for 30 minutes at 4°C. The extracts were centrifuged at 13,000 rpm for 5 minutes at 4°C, after which the supernatants were desalted on a Sephadex G-25 column (Illustra NAP-10 Columns, GE Healthcare, Pittsburgh) and concentrated by rotary evaporation. The concentrated precursors were dissolved in 200 μl HPLC-grade water.

### Peptidoglycan extraction

The peptidoglycan architecture was analyzed as described ^62^. Mycelia of the wild-type strain, *alpha* and the *dcw* mutant complemented with *divIVA* were grown on top of cellophane discs on modified LPMA medium lacking sucrose and horse serum. Following growth, the mycelial mass was removed from the cellophane, washed in 0.1M Tris-HCl pH 7.5 and lyophilized. 10 mg of the lyophilized biomass was used for PG isolation. Therefore, the biomass was boiled in 0.25% SDS in 0.1 M Tris/HCl pH 6.8, thoroughly washed, sonicated, and treated with DNase, RNase and trypsin. Inactivation of these enzymes was performed by boiling the samples followed by washing with water. Wall teichoic acids were removed with 1 M HCl ^63^. PG was digested with mutanolysin and lysozyme. Muropeptides were reduced with sodium borohydride and the pH was adjusted to 3.5-4.5 with phosphoric acid.

### LC-MS analysis of PG precursors and muropeptides

The LC-MS setup consisted of a Waters Acquity UPLC system (Waters, Milford, MA, USA) and an LTQ Orbitrap XL Hybrid Ion Trap-Orbitrap Mass Spectrometer (Thermo Fisher Scientific, Waltham, MA, USA) equipped with an Ion Max electrospray source. Chromatographic separation of muropeptides and precursors was performed on an Acquity UPLC HSS T3 C18 column (1.8 μm, 100 Å, 2.1 × 100 mm). Mobile phase A consisted of 99.9% H_2_O and 0,1% formic acid, while mobile phase B consisted of 95% acetonitrile, 4.9% H_2_O and 0,1% formic acid. All solvents used were of LC-MS grade or better. The flow rate was set to 0.5 ml min^−1^. The binary gradient program consisted of 1 min 98% A, 12 min from 98% A to 85% A, and 2 min from 85% A to 0% A. The column was then flushed for 3 min with 100% B, after which the gradient was set to 98% and the column was equilibrated for 8 min. The column temperature was set to 30°C and the injection volume used was 5 μL. The temperature of the autosampler tray was set to 8°C. Data was collected in the positive ESI mode with a scan range of m/z 500–x2500 in high range mode. The resolution was set to 15.000 (at m/z 400).

### Sequence homology analysis of *dcw* gene clusters

The homology search of the different *dcw* clusters was done using MultiGeneBlast ^64^. The query used for the search was the *dcw* cluster from *Streptomyces coelicolor* A3(2), for which the required sequences were obtained from the *Streptomyces* Annotation Sever (StrepDB). The homology search included the loci from SCO2077 (*divIVA*) until SCO2091 (*ftsL*). A database was constructed with genome assemblies obtained from NCBI. The analyzed species have the following accession numbers: NC_003888 (*S. coelicolor* A3(2), NZ_MPLE00000000.1 (*Kitasatospora viridifaciens* DSM40239), CP000480 (*Mycobacterium smegmatis* MC2 155), AL123456 (*Mycobacterium tuberculosis* H37Rv), CP014279 (*Corynebacterium stationis* ATCC 6872), BX927147 (*Corynebacterium glutamicum* ATCC13032), AL009126 (*Bacillus subtilis subsp.168)*, U00096 (*Escherichia coli* K-12), CP000253.1 (*Staphylococcus aureus* NTC8325), and AE007317 (*Streptococcus pneumoniae* R6). In the homology search, the Blast parameters were set to a minimal sequence coverage of 25% and a minimal identity of 30%. The first 11 hits of the MultiGeneBlast output are shown in Fig. S1, where homologs genes are represented by arrows with the same colors.

### Phylogeny analysis of *Streptomyces* and *Kitasatospora* species

A set of 1050 *Streptomyces* and *Kitasatospora* genomes was downloaded from NCBI by querying the fasta files in combination with the taxonomic identifier. To this set, 116 unpublished draft genome sequences of an in-house collection of actinomycetes were added ^65^. Complete protein sets encoded within the genomes of *Streptomyces* and *Kitasatospora* spp. were extracted. The Pfam domains of four housekeeping proteins, AtpD (ATP synthase subunit beta), RecA (recombinase A), TrpB (tryptophan synthase beta chain) and GyrB (DNA gyrase subunit B), were retrieved from https://pfam.xfam.org/ and are annotated as PF00213, PF00154, PF06233 and PF00204, respectively. Using the selected Pfam domains, the HMMsearch program of the HMMER v3.0 package ^66^ was employed to identify analogous proteins within the chosen species. MAFFT was used to perform a multiple sequence alignment ^67^. Aligned sequences were concatenated using SeqKit ^68^ and maximum likelihood phylogenetic trees were calculated with RAxML ^69^. iTOL ^70^ was used for the visualization of the phylogenetic tree.

### Detection of *murG* genes in *Streptomyces* and *Kitasatospora* species

MurG domains were predicted using the Pfam database ^44^. Proteins with the predicted MurG domains were used to search in the complete protein sets encoded within the extracted genomes using HMMsearch. Instead of a multiple sequence alignment each protein domain sequence was aligned to its profile Hidden Markov model from Pfam using the HMMalign tool ^71^. For each protein a pairwise distance was calculated for all detected MurG proteins and the threshold was set at 0.9. Network visualizations were constructed using Cytoscape (v. 3.7.1) ^72^.

## ACKNOWLEDGEMENTS

We are grateful to Marc Bramkamp for providing us with DivIVA antibodies, and to Eveline Ultee, Joeri Wondergem and Doris Heinrich for help with microscopy. This work was supported by a VIDI grant (12957) from the Dutch Applied Research Council to D.C. and by grant 15812 from NWO-TTW to G.P.v.W. and D.C.

---

**Supplementary Figure 1. Comparative analysis of *dcw* gene clusters from different bacteria.** (A) Organization and content of the *dcw* gene cluster from *Streptomyces coelicolor* A3(2). (B) MultiGeneBlast output showing homologous *dcw* gene clusters with a minimal identity of 30% and minimal sequence coverage of 25% to the *S. coelicolor* cluster.

**Supplementary Figure 2. Localization of DivIVA-eGFP in *alpha*.** (A) Fluorescence microscopy analysis of *alpha* grown in TSBS medium as a mycelium and carrying pKR1 (left panels), pGreen (middle panels) or pKR2 (right panels). In mycelium containing pKR2, localization of DivIVA-eGFP is found at the hyphal tips (see arrowheads in right panels). No fluorescence is observed in mycelium containing the control plasmid pKR1 (left panels), while a cytosolic signal is observed in *alpha* transformed with pGreen (middle panels). (B) Fluorescence microscopy analaysis of *alpha* grown in LPB medium in the wall-deficient state and carrying pKR1 (left panels), pGreen (middle panels) and pKR2 (right panels). Cells expressing the DivIVA-eGFP fusion protein show distinct foci localized to the membrane (right panels). Like in mycelia, no fluorescence is observed in cells containing the control plasmid pKR1 (left panels), while a cytosolic signal is evident in cells containing pGreen (middle panels). Scale bars represent 10 μm.

**Supplementary Figure 3. PCR verification demonstrating the deletions of *divIVA* and the partial *dcw* gene cluster in *alpha*.** (A) Schematic illustration of the *dcw* clusters in *alpha* (top) and the derivative strains lacking *divIVA* (middle) or part of the *dcw* cluster (bottom). To verify the deletions, PCR analyses were performed using primers *divIVA-Fw* and *divIVA-Rv* (B) and *dcw-Fw* and *dcw-Rv* (C). (B) PCR analysis using primers *divIVA-Fw* and *divIVA-Rv* yielded PCR products of 1.8 Kb when chromosomal DNA of the wild-type strain (DSM40239) or *alpha* were used, while a 2.7 Kb fragment was obtained in the Δ*divIVA* mutant. As expected, no product was obtained with these primers using chromosomal DNA of the *dcw* mutant as the template. (C) PCR analysis using primers *dcw-Fw* and *dcw-Rv only* yielded a PCR product of 1.7 Kb when chromosomal DNA of the *dcw* mutant was used as the template. Please note that the sizes of the fragments expected for the wild-type strain and *alpha* (8.2 Kb) and the Δ*divIVA* mutant (9.2 Kb) are too large for efficient amplification.

**Supplementary Figure 4. Domain structure of MurG and MurG2 proteins.** MurG proteins contain an N-terminal domain (PF03033) that binds Lipid I and is involved in membrane association. The C-terminal domain (PF04101) contains the UDP-GlcNAc binding site. These domains are found in MurG proteins of *E. coli* (AAC73201.1), *B. subtilis* (CAB13395.2), *S. coelicolor* (NP_626343.1) and *K. viridifaciens* (BOQ63_RS32465). Notably, MurG2 of *K. viridifaciens* (BOQ63_RS12640) also contains both domains. Please note that the protein encoded by the *BOQ63_RS05415* gene only contains the N-terminal domain (PF03033), but not the C-terminal (PF04101) domain.

**Supplementary Figure 5. Sequence similarity network of the MurG and MurG2 proteins encoded in the genomes of *Streptomyces* and *Kitasatospora* species.** Nodes represent MurG proteins and edges highlight similarity (with a threshold set at 0.9). Node colors indicate if the MurG(-like) proteins are encoded in the *dcw* gene cluster (red) or elsewhere in the genome (green). Circular node shapes are proteins from *Streptomyces* spp., while those from *Kitasatospora* spp. are shown as diamonds. Please note that almost all MurG proteins encoded in the *dcw* cluster group together.

**Supplementary Figure 6. PCR analysis demonstrating the *murG* and *murG2* deletions in *alpha*.** The deletion of *murG* and *murG2* in *alpha* was verified by PCR. In strains carrying a wild-type *murG* gene (DSM40239, *alpha* and Δ*murG2*) a fragment of 1.3 Kb is amplified. In contrast, a fragment of 1.4 Kb is found in *murG* mutants (Δ*murG* and Δ*murG*/Δ*murG2*; left gel). Likewise, the expected PCR product for strains carrying the *murG2* wild-type gene (DSM40239, *alpha*, Δ*murG*) was 1.2 Kb, while replacement of *murG2* by apramycin or viomycin yielded PCR products of 1.3 Kb and 1.5 Kb, respectively (right gel). Please note that the *murG2* gene is still detectable in the Δ*murG*Δ*murG2* merodiploid.

**Supplementary Figure 7. PCR analysis demonstrating the *murG* and *murG2* deletions in *Kitasatospora viridifaciens*.** The deletion of *murG* and *murG2* in *K. viridifaciens* was verified by PCR. In the wild-type strain (DSM40239) a fragment of 1365 bp is amplified, while a fragment of 1436 bp is found in three independent *murG* mutants (Δ*murG*; left gel). Likewise, the expected size of the PCR product for the wild-type strain carrying the *murG2* gene (DSM40239) was 1279 bp, while replacement of *murG2* yielded a PCR product of 1311 bp (Δ*murG2*; right gel).

**Supplementary Movie 1. L-form proliferation of *alpha***. Time-lapse microscopy showing proliferation of *alpha* in LPB medium containing high levels of sucrose. The times are indicated in min. The scale bar indicates 5 μm.

**Supplementary Movie 2. Extrusion of L-forms from hyphal tips.** Cell wall-deficient L-forms are extruded from hyphal tips when mycelium of *alpha* is transferred to LPMA agar containing high levels of sucrose. The times are indicated in min. The scale bar indicates 5 μm.

