## Supplementary Data for "A novel and conserved cell wall enzyme that can substitute for the Lipid II synthase MurG"

Zhang *et al*.

**This PDF file includes:**

Supplementary Figures 1-7
Supplementary Tables 1-4
Supplementary References

### SUPPLEMENTARY FIGURES

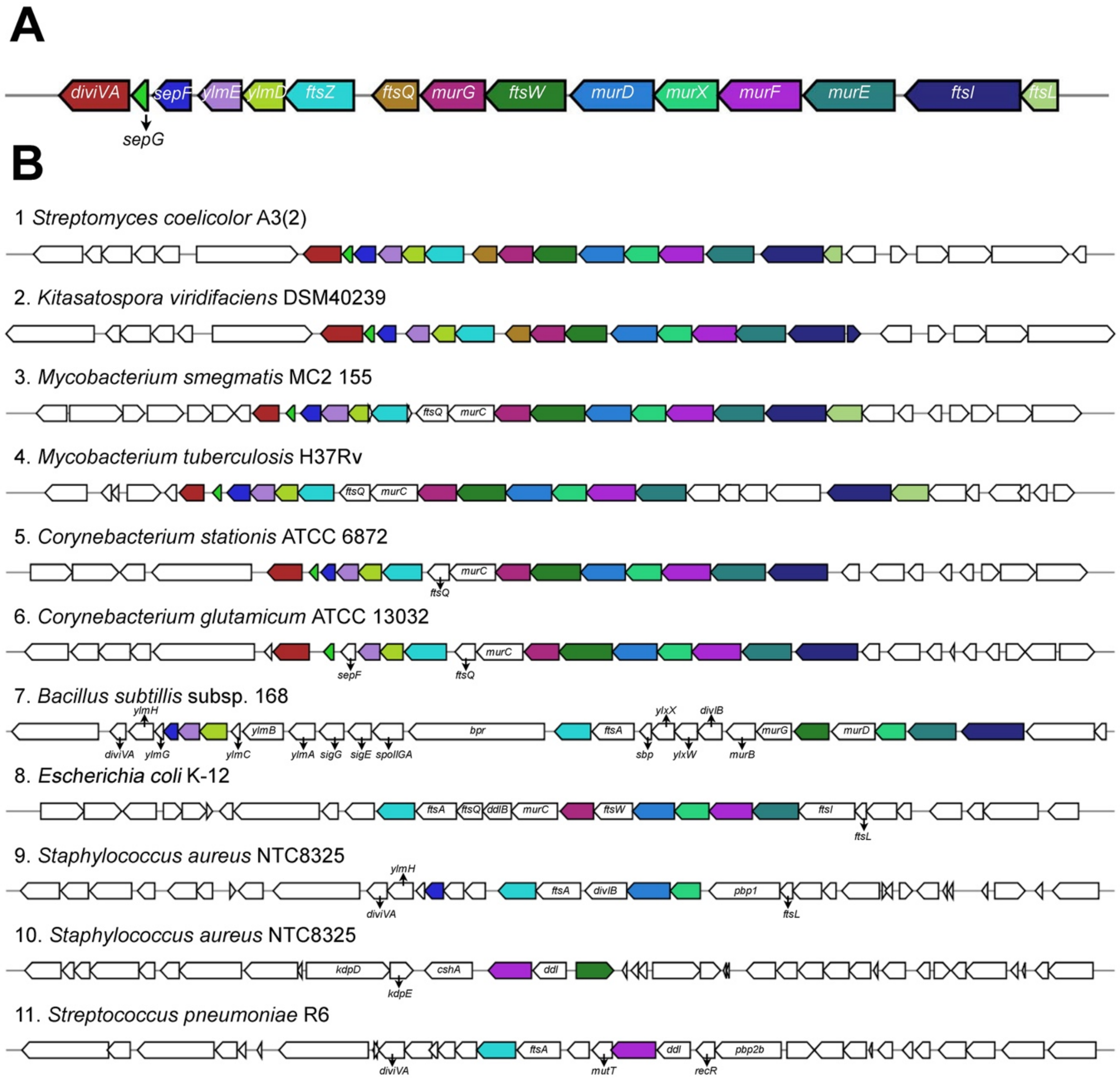


**Supplementary Figure 1. Comparative analysis of *dcw* gene clusters from different bacteria.** (A) Organization and content of the *dcw* gene cluster from *Streptomyces coelicolor* A3(2). (B) MultiGeneBlast output showing homologous *dcw* gene clusters with a minimal identity of 30% and minimal sequence coverage of 25% to the *S. coelicolor* cluster.

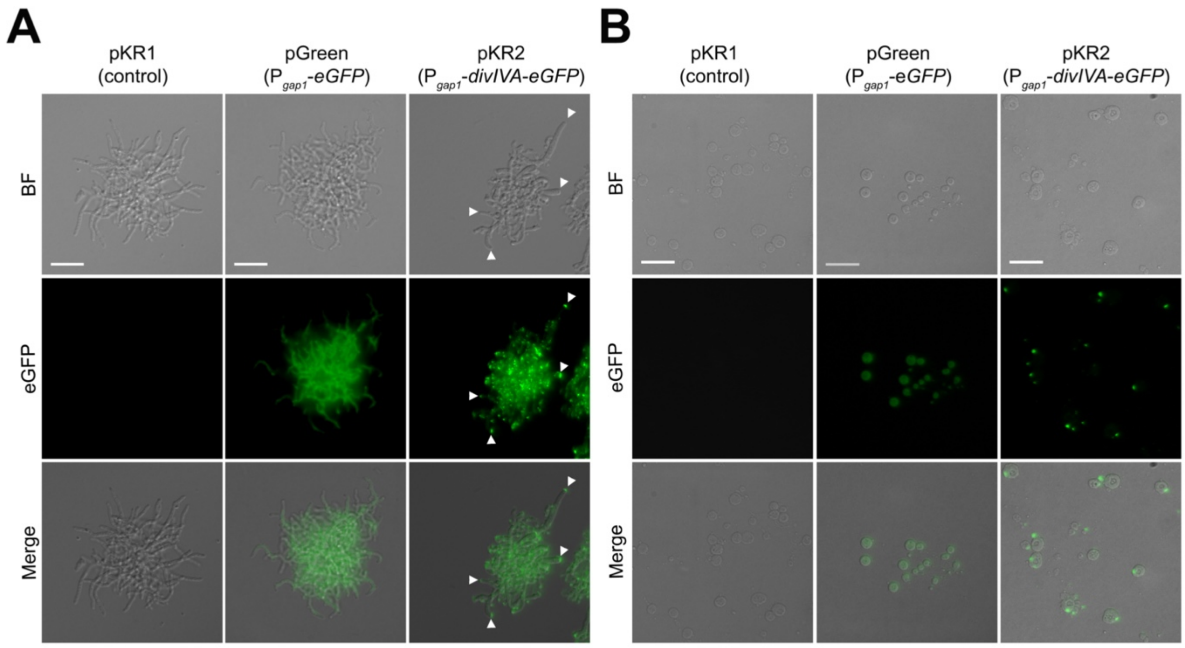


**Supplementary Figure 2. Localization of DivIVA-eGFP in *alpha*.**(A) Fluorescence microscopy analysis of *alpha* grown in TSBS medium as a mycelium and carrying pKR1 (left panels), pGreen (middle panels) or pKR2 (right panels). In mycelium containing pKR2, localization of DivIVA-eGFP is found at the hyphal tips (see arrowheads in right panels). No fluorescence is observed in mycelium containing the control plasmid pKR1 (left panels), while a cytosolic signal is observed in *alpha* transformed with pGreen (middle panels). (B) Fluorescence microscopy analaysis of *alpha* grown in LPB medium in the wall-deficient state and carrying pKR1 (left panels), pGreen (middle panels) and pKR2 (right panels). Cells expressing the DivIVA-eGFP fusion protein show distinct foci localized to the membrane (right panels). Like in mycelia, no fluorescence is observed in cells containing the control plasmid pKR1 (left panels), while a cytosolic signal is evident in cells containing pGreen (middle panels). Scale bars represent 10 µm.

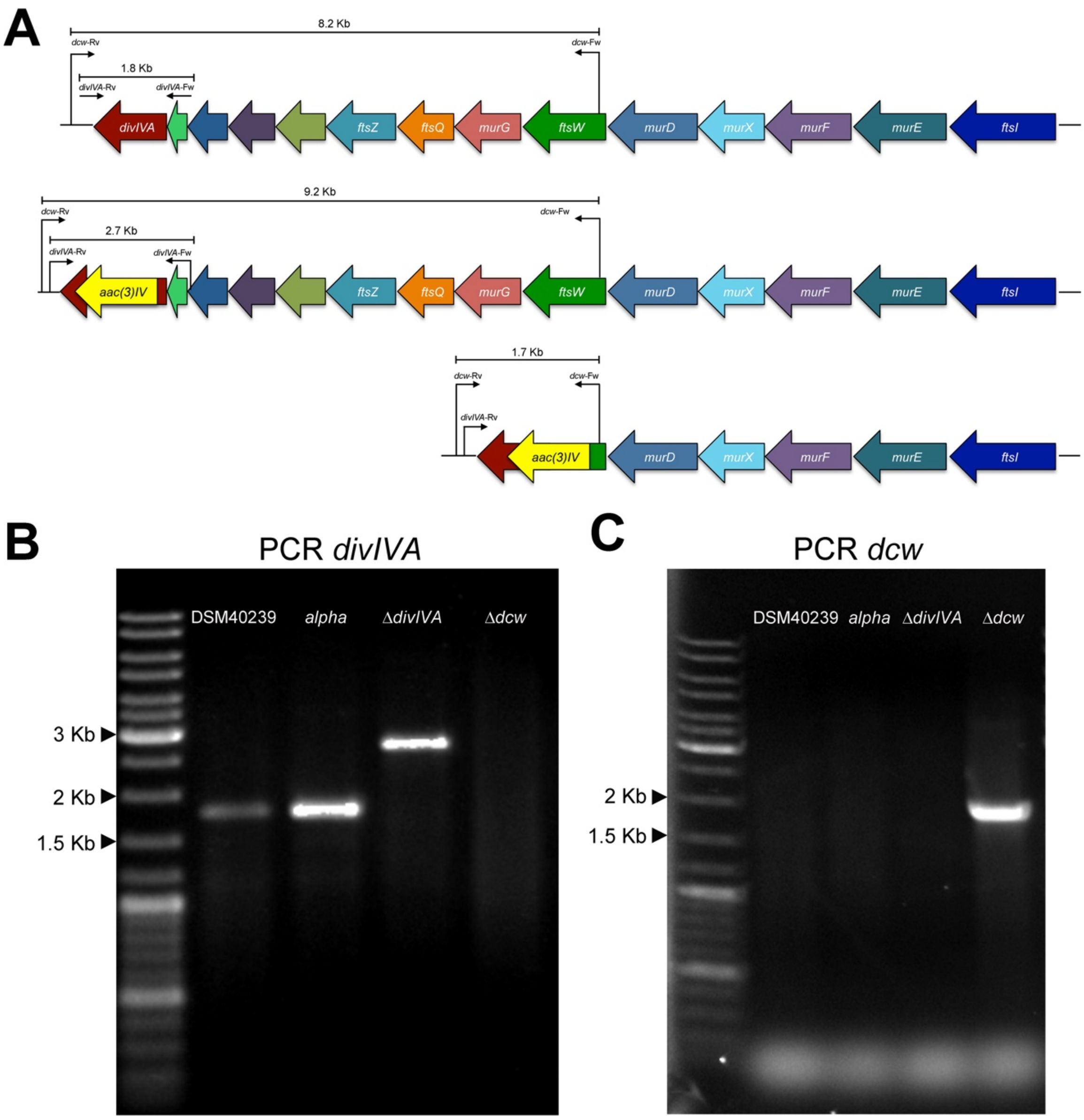


**Supplementary Figure 3. PCR verification demonstrating the deletions of *divIVA* and the partial *dcw* gene cluster in *alpha*.** (A) Schematic illustration of the *dcw* clusters in *alpha* (top) and the derivative strains lacking *divIVA* (middle) or part of the *dcw* cluster (bottom). To verify the deletions, PCR analyses were performed using primers *divIVA-Fw* and *divIVA-Rv* (B) and *dcw-Fw* and *dcw-Rv* (C). (B) PCR analysis using primers *divIVA-Fw* and *divIVA-Rv* yielded PCR products of 1.8 Kb when chromosomal DNA of the wild-type strain (DSM40239) or *alpha* were used, while a 2.7 Kb fragment was obtained in the Δ*divIVA* mutant. As expected, no product was obtained with these primers using chromosomal DNA of the *dcw* mutant as the template. (C) PCR analysis using primers *dcw-Fw* and *dcw-Rv only* yielded a PCR product of 1.7 Kb when chromosomal DNA of the *dcw* mutant was used as the template. Please note that the sizes of the fragments expected for the wild-type strain and *alpha* (8.2 Kb) and the Δ*divIVA* mutant (9.2 Kb) are too large for efficient amplification.

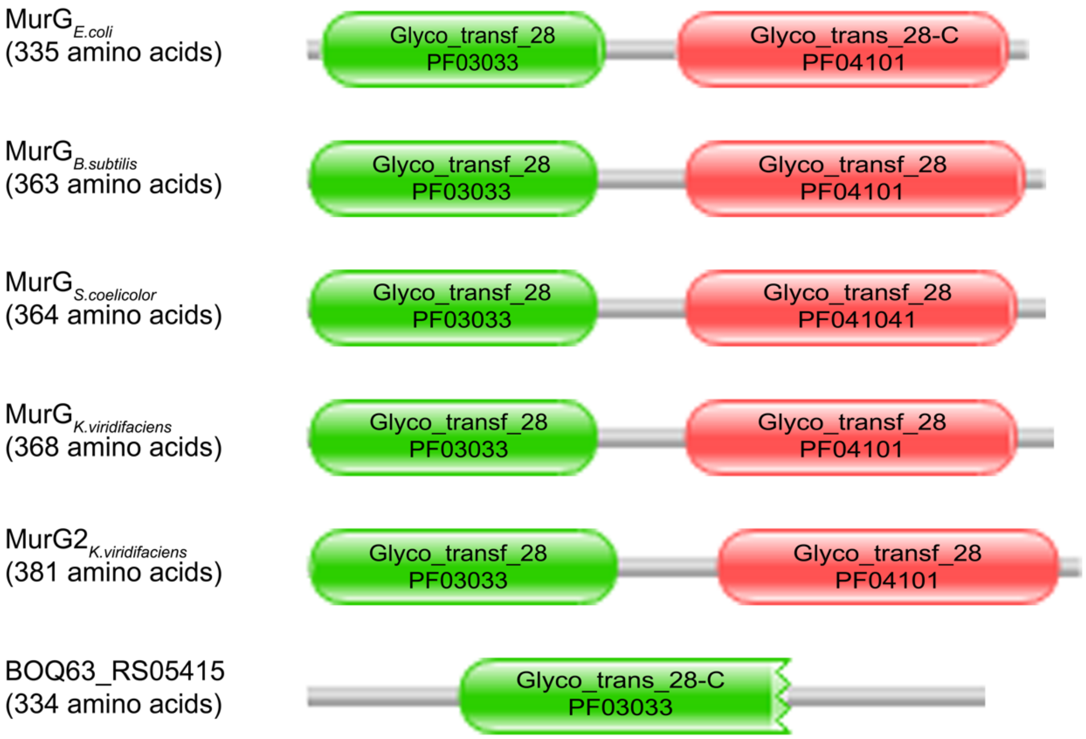


**Supplementary Figure 4. Domain structure of MurG and MurG2 proteins**

MurG proteins contain an N-terminal domain (PF03033) that binds Lipid I and is involved in membrane association. The C-terminal domain (PF04101) contains the UDP-GlcNAc binding site. These domains are found in MurG proteins of *E. coli* (AAC73201.1), *B. subtilis* (CAB13395.2), *S. coelicolor* (NP_626343.1) and *K. viridifaciens* (BOQ63_RS32465). Notably, MurG2 of *K. viridifaciens* (BOQ63_RS12640) also contains both domains. Please note that the protein encoded by the *BOQ63_RS05415* gene only contains the N-terminal domain (PF03033), but not the C-terminal (PF04101) domain.

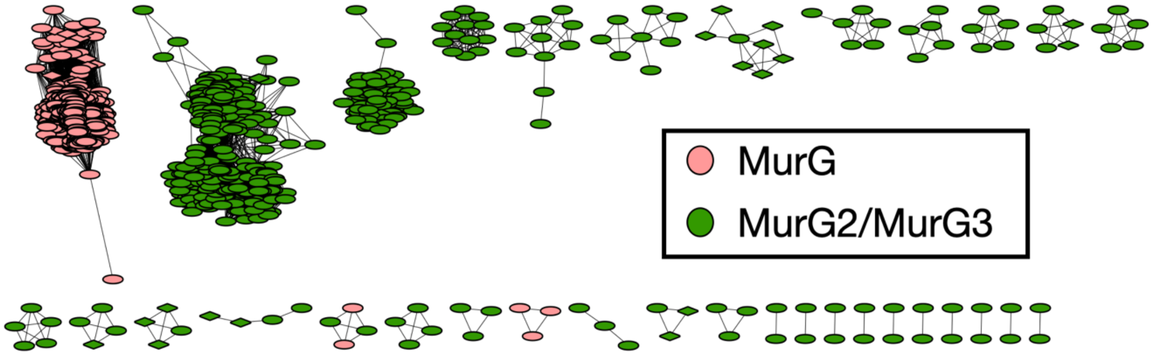


**Supplementary Figure 5. Sequence similarity network of the MurG and MurG2 proteins encoded in the genomes of *Streptomyces* and *Kitasatospora* species.** Nodes represent MurG proteins and edges highlight similarity (with a threshold set at 0.9). Node colors indicate if the MurG(-like) proteins are encoded in the *dcw* gene cluster (red) or elsewhere in the genome (green). Circular node shapes are proteins from *Streptomyces* spp., while those from *Kitasatospora* spp. are shown as diamonds. Please note that almost all MurG proteins encoded in the *dcw* cluster group together.

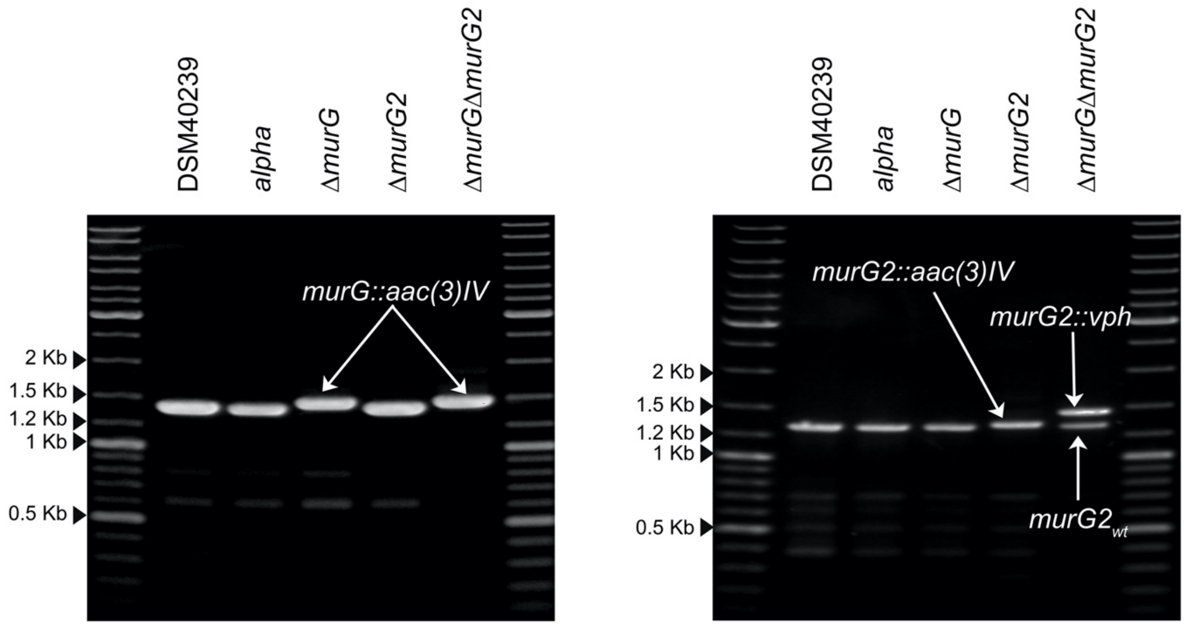


**Supplementary Figure 6. PCR analysis demonstrating the *murG* and *murG2* deletions in *alpha*.** The deletion of *murG* and *murG2* in *alpha* was verified by PCR. In strains carrying a wild-type *murG* gene (DSM40239, *alpha* and Δ*murG2)* a fragment of 1.3 Kb is amplified. In contrast, a fragment of 1.4 Kb is found in *murG* mutants (Δ*murG* and Δ*murG/*Δ*murG2*; left gel). Likewise, the expected PCR product for strains carrying the *murG2* wild-type gene (DSM40239, *alpha*, Δ*murG*) was 1.2 Kb, while replacement of *murG2* by apramycin or viomycin yielded PCR products of 1.3 Kb and 1.5 Kb, respectively (right gel). Please note that the *murG2* gene is still detectable in the Δ*murG*Δ*murG2* merodiploid.

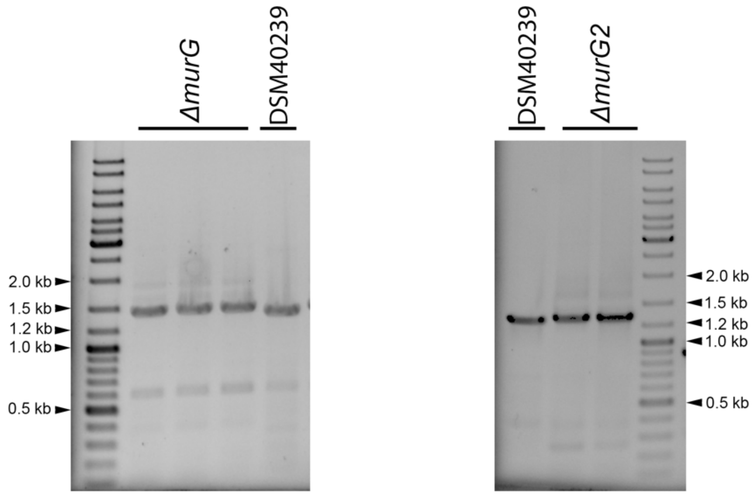


**Supplementary Figure 7. PCR analysis demonstrating the *murG* and *murG2* deletions in *Kitasatospora viridifaciens.*** The deletion of *murG* and *murG2* in *K. viridifaciens* was verified by PCR. In the wild-type strain (DSM40239) a fragment of 1365 bp is amplified, while a fragment of 1436 bp is found in three independent *murG* mutants (Δ*murG*; left gel). Likewise, the expected size of the PCR product for the wild-type strain carrying the *murG2* gene (DSM40239) was 1279 bp, while replacement of *murG2* yielded a PCR product of 1311 bp (Δ*murG2*; right gel).

### SUPPLEMENTARY TABLES

**Supplementary Table 1.** Strains used in this study

| Strains | Genotype | Reference |
|---|---|---|
| ***E. coli* strains** | | |
| DH5α | F- Φ80lacZDM15 D(lacZYA-argF)U169 recA1 endA1 hsdR17(rK-, mK-) phoA supE44 thi-1 gyrA96 relA1 λ- | 1 |
| JM109 | *recA1, endAl*, *gyrA*96, *thi, hsdR*17, *supE*44, *relA*1, $\lambda^-$, Δ(*lac-proAB*), [F', *traD*36, *proAB*, Δ(lacI$^q$ZΔM15] | 2 |
| ET12567 | F-*dam-13::*Tn9 *dcm-6 hsdM hsdR recF143 zjj-202::*Tn10 *galK2 galT22 ara14 lacY1 xyl-5 leuB6 thi-1 tonA31 rpsL136 hisG4 tsx78 mtl-1 glnV44* | 3 |
| SCS110 | *rpsL* (Str$^r$) *thr leu endA thi-1 lacY galK galT ara tonA tsx dam dcm supE*44 Δ(*lac-proAB*) [*F´ traD*36 *proAB lacI$^q$* ZΔM15] | 4 |
| **Actinobacteria** | | |
| *Streptomyces coelicolor* A3(2) M145 | Wild-type strain | Lab collection |
| M145 + pGWS1379 | *S. coelicolor* A3(2) M145 expressing *murG2* | This work |
| *Kitasatospora viridifaciens* DSM40239 | Wild-type strain | DSMZ[5] |
| DSM40239Δ*murG* | *K. viridifaciens* DSM40239 in which *murG* is replaced by the *aac(3)IV* apramycin resistance cassette | This work |
| DSM40239Δ*murG2* | *K. viridifaciens* DSM40239 in which *murG2* is replaced by the *aac(3)IV* apramycin resistance cassette | This work |
| ***K. viridifaciens* L-form strains** | | |
| *alpha* | L-form cell line obtained after induction with penicillin and lysozyme | 6 |
| *alpha+pKR1* | *alpha* carrying pKR1 | This work |
| *alpha+pKR2* | *alpha* carrying pKR2 | This work |
| *alpha+pGreen* | *alpha* constitutively expressing *eGFP* | This work |
| Δ*divIVA* | *alpha* in which *divIVA* is replaced by the *aac(3)IV* apramycin resistance cassette | This work |
| Δ*dcw* | *alpha* in which *ftsW, murG, ftsQ, ftsZ, ylmD, ylmE, sepG, sepF* and *divIVA* are replaced by the *aac(3)IV* apramycin resistance cassette | This work |
| Δ*divIVA+divIVA* | *divIVA* mutant containing *divIVA* expressed from the *gap1* promoter | This work |
| Δ*dcw+divIVA* | *dcw* mutant containing *divIVA* expressed from the *gap1* promoter | This work |
| Δ*murG* | *alpha* in which *murG* is replaced by the *aac(3)IV* apramycin resistance cassette | This work |
| Δ*murG2* | *alpha* in which *murG2* is replaced by the *aac(3)IV* apramycin resistance cassette | This work |
| Δ*murG*Δ*murG2* | Δ*murG1* in which *murG2* is replaced by the *vph* viomycin resistance cassette | This work |

**Supplementary Table 2.** Vectors and constructs used in this study

| Plasmid | Description and relevant features | Reference |
|---|---|---|
| pWHM3 | Unstable, multi-copy and self-replicating S*treptomyces* vector. Contains thiostrepton and ampicillin resistance cassette. | 7 |
| pIJ780 | Plasmid containing a viomycin (*vph*) resistance cassette. | 8 |
| pIJ8600 | *E. coli–Streptomyces* shuttle vector containing the φC31 *attP-int* region for genomic integration. Confers resistance to apramycin and thiostrepton. | 9 |
| pIJ8630 | *E. coli–Streptomyces* shuttle vector containing the φC31 *attP-int* region for genomic integration. Confers resistance to apramycin | 9 |
| pSET152 | *E. coli/Streptomyces* shuttle vector, high copy number in *E. coli* and integrative in *Streptomyces* | 10 |
| pHM10a | Conjugative *E. coli-Streptomyces* shuttle vector, harboring P*ermE* and RBS | 11 |
| pMS82 | *E. coli/Streptomyces* shuttle vector, high copy number in *E. coli* and integrative in *Streptomyces* | 12 |
| pGreen | pIJ8630 containing the *eGFP* gene under control of the constitutive *gap1* promoter of *S. coelicolor.* | 13 |
| pKR1 | pIJ8630 derivative containing the viomycin resistance cassette from pIJ780 cloned into the unique NheI site. | This work |
| pKR2 | pKR1 derivative containing a C-terminal *eGFP* fusion to *divIVA* of *K. viridifaciens* under control of the *gap1* promoter of *S. coelicolor.* | This work |
| pKR3 | pWHM3 containing the flanking regions of the *K. viridifaciens divIVA* gene interspersed by the *apra-loxP* cassette. | This work |
| pKR4 | pWHM3 derivative containing the flanking regions around the *K. viridifaciens* partial *dcw* gene cluster (*ftsW*, *murG*, *ftsQ*, *ftsZ*, *ylmD*, *ylmE*, *sepF*, *sepG*, *divIVA*) interspersed by the *apra-loxP* cassette. | This work |
| pKR5 | pIJ8600 derivative containing the *gap1* promoter of *S. coelicolor*. | This work |
| pKR6 | pKR5 derivative containing the *divIVA* gene of *K. viridifaciens* under control of the *gap1* promoter of *S. coelicolor*. | This work |
| pKR8 | pWHM3 containing the flanking regions of the *K. viridifaciens murG* gene interspersed by the *apra-loxP* cassette. | This work |
| pKR9 | pWHM3 containing the flanking regions of the *K. viridifaciens murG2* (BOQ63_RS12640) gene interspersed by the *apra-loxP* cassette. | This work |
| pKR10 | pWHM3 containing the flanking regions of the *K. viridifaciens murG2* (BOQ63_RS12640) gene interspersed by the viomycin resistance cassette. | This work |
| pGWS1369 | pSET152 lacking its NcoI site | This work |
| pGWS1370 | pGWS1369 containing sgRNA scaffold (no spacer) and P*gapdh-dCas9* | This work |
| pGWS1371 | pWGS1370 containing spacer targeting the template strand of *SCO2084* | This work |
| pGWS1376 | pWGS1370 containing spacer targeting the non-template strand of *SCO2084* | This work |
| pGWS1378 | pSET152 containing P*ermE-murG2* | This work |
| pGWS1379 | pMS82 containing P*ermE-murG2* | This work |

**Supplementary Table 3.** Primers used in this study

| Primer | Sequence (5' – 3') |
|---|---|
| *vph*-FW-NheI | GACGCTAGCGGCTGACGCCGTTGGATACACCAAG |
| *vph*-RV-NheI | GACGCTAGCAATCGACTGGCGAGCGGCATCCTAC |
| $P_{Gap1}$-FW-BglII | GATTACAGATCTCCGAGGGCTTCGAGACC |
| $P_{Gap1}$-RV-XbaI | GATGACTCTAGACCGATCTCCTCGTTGGTAC |
| *divIVA*-FW-XbaI | GTCAAGCTTCTAGAATGCCATTGACCCCCGAGGA |
| *divIVA*-Nostop-RV-NdeI | GACCATATGGTTGTCGCCGTCCTCGTCAATCAGG |
| P1-*divIVA*-FW | GACGACGAATTCTGTGATGACCGTCGCTCCACTG |
| P2-*divIVA*-RV | GACGACTCTAGACTTCCGCATGTTGGCCTGGTTC |
| P1-*dcw*-FW | GACGAATTCTCCGCGAGGTCACGTACATC |
| P2-*dcw*-RV | GACTCTAGAAGAGCACCAGTGCGAGCTTG |
| P3-*dcw*-FW | GACTCTAGAAGCAGCAGATGGGCAACCAG |
| P4-*dcw*-RV | GATAAGCTTCCCGGCTACAACCTCAGTTGTC |
| Delcheck-*divIVA*-FW | TGACCCGGCCACGACTTTAC |
| Delcheck-*divIVA*-RV | GGACGCCCTCAACAAAC |
| Delcheck-*dcw*-FW | CCAGAACTGGCTGGATTTCG |
| Delcheck-*dcw*-RV | GTCTCCAGGTACGACTTCAG |
| *divIVA*-XbaI-FW | GTCAAGCTTCTAGAATGCCATTGACCCCCGAGGA |
| *divIVA*-NdeI-RV | GATCGAATTCATATGCCCGGCTACAACCTCAGTTGTC |
| *divIVA* seq1-FW | AGCAGCAGATGGGCAACCAG |
| *divIVA* seq2-FW | CGCGTCTGAAGTCGTACCTG |
| *divIVA* seq-RV | ACCTCGTCCTCGTCATAGC |
| SCO2079_F-520 | TCACGGCGCTGTCGAAGGAGGCCG |
| SCO2079_R+1162 | CTCATCGAGGAAGGCATCGACCTC |
| *divIVA*$_{SCO}$-FW | AAGGCTACGCCGTACTACAG |
| *divIVA*$_{SCO}$-RV | AGATACGGGCTTGCCGAATG |
| P1-*murG*-Fw | CATCGAATTCGATATCTTTCGGCTTCTTCCAGTTCC |
| P2-*murG*-Rv | CATCCATGTCTAGACGACATGCACCGAAATTCAC |
| P3-*murG*-Fw | CATCCATGTCTAGATGGTGTACGAGGCGATCCAG |
| P4-*murG*-Rv | CATGGATATCAAGCTTGACGGATGTCGATGGGTAGG |
| Delcheck-*murG*-Fw | AGCAAGAACTCCCGGATCAG |
| Delcheck-*murG*-Rv | AGCACCGACGAGAAGAACAC |
| P1-*mglB*-Fw | CTGAGAATTCGATATCTTCTCGTGGGAACACCGGGCA |
| P2-*mglB*-Rv | CTGATCTAGAGGTGACGATCAGCCGCATAGG |
| P3-*mglB*-Fw | CTGATCTAGAGACCGTCTCGTGGACGTGCTG |
| P4-*mglB*-Rv | CTGAAAGCTTGATATCGTTCCCGCTACCCGAACGGAAC |
| Delcheck-*murG2*-Fw | CTGAATGTTCCAAGCGTGAACCGGGA |
| Delcheck-*murG2*-Rv | CTGAGCGACTACAAGGCGTACCAGG |
| *vph*-Fw-EcoRI-HindIII-XbaI | GACGAATTCAAGCTTTCTAGAGGCTGACGCCGTTGGATACACCAAG |
| *vph*-Rv-EcoRI-HindIII-XbaI | GACGAATTCAAGCTTTCTAGAAATCGACTGGCGAGCGGCATCCTAC |
| 152DNcoI_F | GCAAGCCATTCTGTCCGCGATGGACAAGCTGTACT |
| 152DNcoI_R | GCAGTACAGCTTGTCCATCGCGGACAGAATGGCTT |
| SgTermi_R_B | ctagGGATCCCAAAAAACCCCTCAAGACCCGTTTAGAGGCCCCAAGG GGTTATGCTAGTTACGCCTACGTAAAAAAAGCACCGACTCGGTGCC |
| SCO2084_T_F | CATGCCATGGACCGTGGGGATCACGGCCCTGTTTTAGAGCTAGAA ATAGC |
| SCO2084_NT5_F | CATGCCATGGTTGGCCTCGTGCACGACGATGTTTTAGAGCTAGAAA TAGC |
| murG2_F+4_ENdeI | ctgaGAATTCCATATGCGGCTGATCGTCACCGGCG |
| murG2_R+1146_HX | ctgaAAGCTTTCTAGACTAGCGGTCCACTACCGACAGCAGCAC |

**Supplementary Table 4.** *murG* homologs in *Kitasatospora viridifaciens*

| Hit | Scaffold | Hit start | Hit end | Locus | Pairwise identity (%) |
|---|---|---|---|---|---|
| 1 | Chromosome | 5,334,877 | 5,335,956 | BOQ63_RS32465 | 100 |
| 2 | Chromosome | 1,072,546 | 1,073,598 | BOQ63_RS12640 | 31.2 |
| 3 | KVP1 | 1,258,806 | 1,257,943 | BOQ63_RS05415 | 16.5 |
